# Balancing doses of EL222 and light improves optogenetic induction of protein production in *Komagataella phaffii*

**DOI:** 10.1101/2024.12.31.630935

**Authors:** Shannon M. Hoffman, Sebastián Espinel-Ríos, Makoto A. Lalwani, Sarah K. Kwartler, José L. Avalos

## Abstract

*Komagataella phaffii*, also known as *Pichia pastoris*, is a powerful host for recombinant protein production, in part due to its exceptionally strong and tightly controlled P_AOX1_ promoter. Most *K. phaffii* bioprocesses for recombinant protein production rely on P_AOX1_ to achieve dynamic control in two-phase processes. Cells are first grown under conditions that repress P_AOX1_ (growth phase), followed by methanol-induced recombinant protein expression (production phase). In this study, we propose a methanol-free approach for dynamic metabolic control in *K. phaffii* using optogenetics, which can help enhance input tunability and flexibility in process optimization and control. The light-responsive transcription factor EL222 from *Erythrobacter litoralis* is used to regulate protein production from the P_C120_ promoter in *K. phaffii* with blue light. We used two system designs to explore the advantages and disadvantages of coupling or decoupling EL222 integration with that of the gene of interest. We investigate the relationship between EL222 gene copy number and light dosage to improve production efficiency for intracellular and secreted proteins. Experiments in lab-scale bioreactors demonstrate the feasibility of the outlined optogenetic systems as potential alternatives to conventional methanol-inducible bioprocesses using *K. phaffii*.

## Introduction

*Komagataella phaffii*, also known as *Pichia pastoris*, is a powerful host for recombinant protein production across the food, pharmaceutical, and enzyme industries^1^. This yeast’s key strengths include its ability to grow to high cell densities, efficiently secrete recombinant proteins, thrive in simple culture conditions compared to mammalian cells, and execute eukaryotic post-translational modifications like disulfide bonds and glycosylation. Disulfide bond formation in the endoplasmic reticulum is relatively simpler than in *Escherichia coli*, which requires targeting the protein to the periplasm or producing it as inclusion bodies and then solubilizing and refolding *in vitro*^2–4^. Even relative to other yeast workhorses like *Saccharomyces cerevisiae, K. phaffii*’s post-translational modifications for glycosylated proteins can be particularly desirable, as *K. phaffii* does not hyperglycosylate^5^. In addition, *K. phaffii* is equipped with strong and tightly-controlled promoters like P_AOX1_^5,6^, which is widely used for protein expression across industries.

The P_AOX1_ promoter’s inducible nature enables *K. phaffii* bioprocesses to operate in two temporal phases, addressing the inherent competition between growth and production. During an initial growth phase, cells are fed glycerol or glucose, allowing them to rapidly grow while P_AOX1_, controlling the recombinant proteins, is repressed. This avoids the metabolic burden and potential toxicity of the recombinant protein to promote rapid cell growth. Once the desired cell density is reached, the production phase is initiated by switching the carbon source to methanol, which strongly induces P_AOX1_ to express the recombinant protein of interest. While effective, using methanol, like other chemical inducers of gene expression, imposes challenges due to its limited tunability and irreversibility. Once methanol is added to the medium, (corrective) input actions for process optimization and control are limited by the difficulty of dynamically titrating methanol concentrations in real-time and lack of a tunable promoter response to methanol. Furthermore, methanol is often produced from unsustainable, non-renewable sources such as coal, natural gas, or petroleum, despite noteworthy ongoing efforts to produce methanol more sustainably, for example, from CO_2_ ^7,8^. Methanol also poses flammability and toxicity hazards, and its utilization in *K. phaffii* processes has high oxygen demand and heat evolution, which constrains reactor size^11^. The similarly strong P_GAP_ promoter offers a methanol-free alternative for high protein expression in glucose; however, its constitutive nature makes it suboptimal for toxic proteins and other targets that benefit from dynamic control in two-phase processes. Thus, for *K. phaffii* bioprocesses to reach their full potential for recombinant protein production, the development of competitive methanol-free inducible systems is of great interest.

Several studies in recent years have focused on developing new methanol-free inducible systems in *K. phaffii*. These efforts involve either altering the regulation of P_AOX1_ or inducing production from alternative promoters. For example, P_AOX1_ expression in glycerol has been achieved up to 77% of wild-type methanol induction by knocking out the endogenous repressors (Δmig1Δmig2Δnrg1) while overexpressing the Mit1 activator^12^. Other studies have similarly reported derepression of P_AOX1_ in glycerol by upregulating activators^13,14^ and knocking out relevant kinases^15^. In these studies, P_AOX1_ remains repressed in glucose, making these strains useful for dynamic processes consisting of a glucose-fed growth phase and a glycerol-fed production phase. While these approaches maintain the tight control of P_AOX1_, they rarely match the strength of methanol-induction of P_AOX1_. Additional studies have turned toward derepressed expression from other promoters involved in methanol utilization like P_DAS1_^16^, P_PDF_^17^, P_FMD_^18^ from *K. phaffii* as well as P_HpMOX_ and P_HpFMD_ from *Hansenula polymorpha*^19^; however, direct comparison of these promoters to P_AOX1_ has either not been reported (P_DAS1_, P_PDF_, P_FMD1_) or not matched the strength of P_AOX1_ (P_HpMOX_, P_HpFMD_). Perhaps the most popular alternative in the methanol utilization pathway is the P_FLD1_ promoter, which is of comparable strength to P_AOX1_ and can be induced using methylamine as a nitrogen source^20–23^. Though effective, this inducer is more expensive than the typically used ammonium sulfate, making it suboptimal for scaleup. Finally, promoters regulated by alternative stimuli like engineered forms of P_ADH2_ (ethanol)^24^, P_THI11_ (thiamine)^25^, P_DH_ (carbon-starvation)^26^, and an engineered variant of P_GTH1_ (glucose concentration)^27^ have been explored, but in many cases (with the exception of P_GTH1_) they suffer from long fed-batch times or weaker expression than P_AOX1_.

In this work, we propose a methanol-free, optogenetic induction approach that modulates gene expression with light rather than with changes in media composition. Optogenetic systems use light-responsive proteins, often transcription factors, to initiate gene expression in response to specific wavelengths of light^28^. In addition to eliminating the need for chemical inducers or media changes, light can be easily applied and removed at varying intensities and time schedules, providing tunability, reversibility, and real-time modulation. Optogenetics has been applied with great success in various microbial hosts to regulate chemical and protein production^29–38^, with several tools based on the transcription factor EL222 from *Erythrobacter litoralis*. EL222 exists as inactive monomers in the dark, which dimerize in blue light (optimally 450 nm) to bind the P_C120_ promoter and activate transcription^39^. In a recent study, EL222 was shown to be capable of inducing gene expression in *K. phaffii* with light^40^, although the optogenetic production of intracellular protein was approximately half that of P_AOX1_, and there was no reported comparison between light and methanol induction for secreted protein production.

Here, we investigate the interplay between EL222 copy number and light dose in protein induction (Fig. 1), offering additional degrees of freedom for process design, optimization, and control. Two strain design strategies are outlined, differing in whether integration of EL222 is coupled with the gene of interest, offering varying flexibility in modulating the copy numbers of these elements. We propose a hybrid modeling framework with Gaussian-process-predicted parameters that can link EL222 copy number and light dose (*manipulated variables*) to the dynamic behavior of the process. We explore the use of EL222 in *K. phaffii* to produce optogenetically regulated intracellular and secreted proteins up to lab-scale bioreactor level, providing new insights into process optimization. As a proof of concept, we compare the proposed optogenetic approach with the more established methanol-inducible system across various cases in recognition of *K. phaffii’*s versatile applications for food, enzyme, material, and pharmaceutical production. Even though the optogenetic systems generally outperform their methanol-based counterparts, under the tested conditions, it is important to note that the aim of this study is not to conclusively determine whether the optogenetic approach is superior. Rather, this depends on the specific case and context regarding multiple technical, design, safety, and economic factors. Instead, we aim to demonstrate the potential of light as a flexible and tunable degree of freedom in recombinant protein production by *K. phaffii*, and to encourage the bioprocess and metabolic engineering communities to add the outlined methods to their toolbox for process design and optimization.

**Figure 1.**
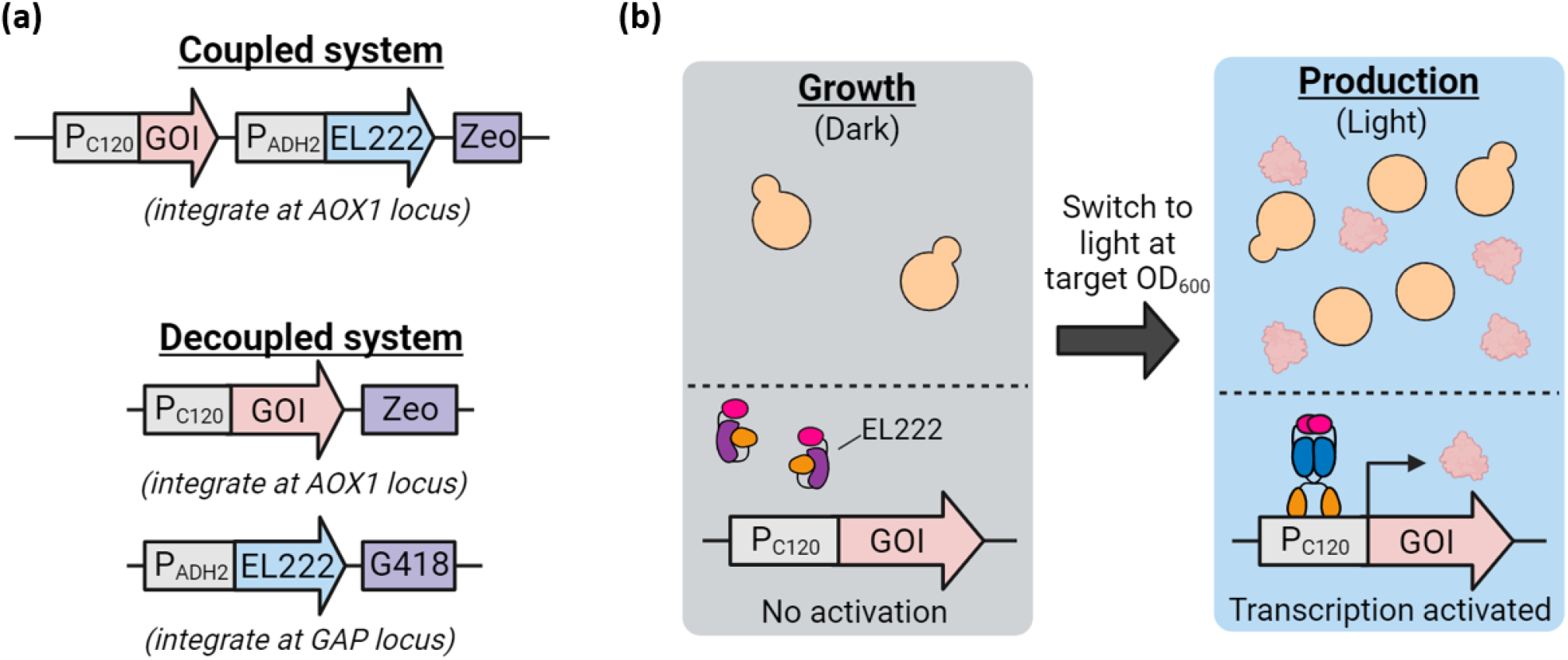
Coupled and decoupled optogenetic systems to induce protein production with light. **(a)** The ‘coupled’ system (top panel) integrates EL222 and the gene of interest together on the same DNA fragment, while the ‘decoupled’ system (bottom panel) integrates these components separately, allowing for different copy numbers of EL222 and of the gene of interest. **(b)** In both cases, the process can be divided into a growth phase in the dark (in which the P_C120_ promoter is inactive) followed by a light-induced production phase, in which EL222 dimerizes upon blue light stimulation (465 nm) to activate transcription from the P_C120_ promoter. Abbreviations: GOI-gene of interest, Zeo-zeocin resistance marker.

## Materials and Methods

### Plasmid construction

All plasmids were cloned into DH5α *E. coli* cells prepared using the Inoue method^71^. We assembled these constructs using Gibson assembly^72^ or restriction ligation with enzymes purchased from NEB. Primers were purchased from Integrated DNA Technologies, and fragments were PCR amplified using CloneAmp Hifi PCR premix from Takara Bio. The Mfp5, SR18, and β-lactoglobulin genes were codon-optimized for expression in *K. phaffii* and synthesized by Twist Bioscience, while other sequences, including those of the P_C120_ promoter and EL222 (Supplemental Sequences S1 and S2) were obtained from lab plasmids or genomic *K. phaffii* DNA. Plasmids and fragments were extracted and purified using Epoch DNA Miniprep and Omega E.Z.N.A. Gel Extraction kits. Final constructs were sequenced using Sanger sequencing by Genewiz or nanopore sequencing through Plasmidsaurus. Detailed descriptions of all plasmids constructed in this study can be found in Supplementary Table S1.

### Yeast strains and transformations

All yeast strains were derived from NRRL Y-11430, which was acquired from ATCC. We carried out transformations using the standard condensed electroporation method^73^, and the resulting strains are catalogued in Supplementary Table S2. Transformants were selected with either zeocin or G418 at concentrations ranging from 100-1000 μg/mL and incubated for 2-3 days at 30 °C until colonies formed. All experimental growth was performed at 30 ºC.

### Quantification of copy numbers by qPCR

Copy numbers of yeast-enhanced green fluorescent protein (yEGFP) (Supplementary Fig. S1) and EL222 (Supplementary Fig. S2 and S3) were quantified by qPCR as previously described by Abad^74^. We extracted genomic DNA from the yeast using phenol-chloroform^75^ and further purified it to remove RNA with RNase A (1 μL of a 20 mg/mL stock per sample)^76^. The qPCR reactions were run using SYBR Green qPCR Master Mix (Thermo Fisher) in a ViiA 7 Real-Time PCR System (Thermo Fisher) with a 96-well adapter. Threshold values were selected by the QuantStudio software (Thermo Fisher), and we used *ARG4* as a housekeeping reference gene to control for any variations in DNA concentration between samples. Primers for amplifying yEGFP, EL222, and *ARG4* were purchased from Integrated DNA Technologies and can be found in Supplementary Table S3. A strain containing a single copy of yEGFP and EL222 (ySMH3-100-3) was identified as described by Abad^74^ using two colony PCRs to check for tandem integrated copies and used as the control from which calibration curves were produced. To identify strains of varying copy numbers, strains were selected from plates ranging in zeocin or G418 concentration from 100-1000 μg/mL. The antibiotic concentration is indicated by the second number in the ySMH3 and ySMH5 strains (for example, the ySMH3-100-3 strain was selected on a 100 μg/mL zeocin plate).

### Illumination and measurement of light intensity

For all microplate experiments, blue (465 nm) light was applied using LED panels (HQRP New Square 12” Grow Light Blue LED 14W) placed above the culture plate. The distance was adjusted for each panel (between 40-70 cm) to achieve the desired light intensity for each experiment (between 5 and 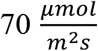), which was measured using a Quantum meter (Apogee Instruments, Model MQ-510). Cultures grown in the dark were shielded from light using aluminum foil.

Since the caps and holding clamps for Erlenmeyer flasks would obstruct light and complicate application of a consistent intensity, light was applied using a different approach for flask experiments. Instead of illuminating from above, an apparatus was constructed to allow illumination by an LED panel from below (Supplementary Fig. S4). For this setup, we secured a light panel to the shaker with four brackets, facilitating the exchange of panels to achieve different light intensities. The panel was covered with a 12" x 12" x ⅛" frosted acrylic sheet (AZM Displays APSheet1/8Milky) with a reusable transparent adhesive pad on top (Chemglass CLS-4010). As with the microplate experiments, intensity was measured at 465 nm with a Quantum meter (Apogee Instruments, Model MQ-510).

### Intracellular production of yEGFP using the coupled optogenetic system and P_AOX1_

Experiments comparing production of intracellular yEGFP by the coupled optogenetic system and P_AOX1_ were performed in both low and high copy strains (Fig. 2, Supplementary Fig. S5 and S6). The optogenetic strains were obtained by transforming the NRRL Y-11430 wild-type strain with a linearized plasmid (using PmeI) containing both P_ADH2_-EL222 and P_C120_-yEGFP (SMH131). Methanol-induced strains were created by transforming NRRL Y-11430 with a similarly linearized construct containing P_AOX1_-yEGFP (SMH5). Transformants were plated on 100 and 1000 μg/mL zeocin to select for both low- and high-copy strains.

**Figure 2.**
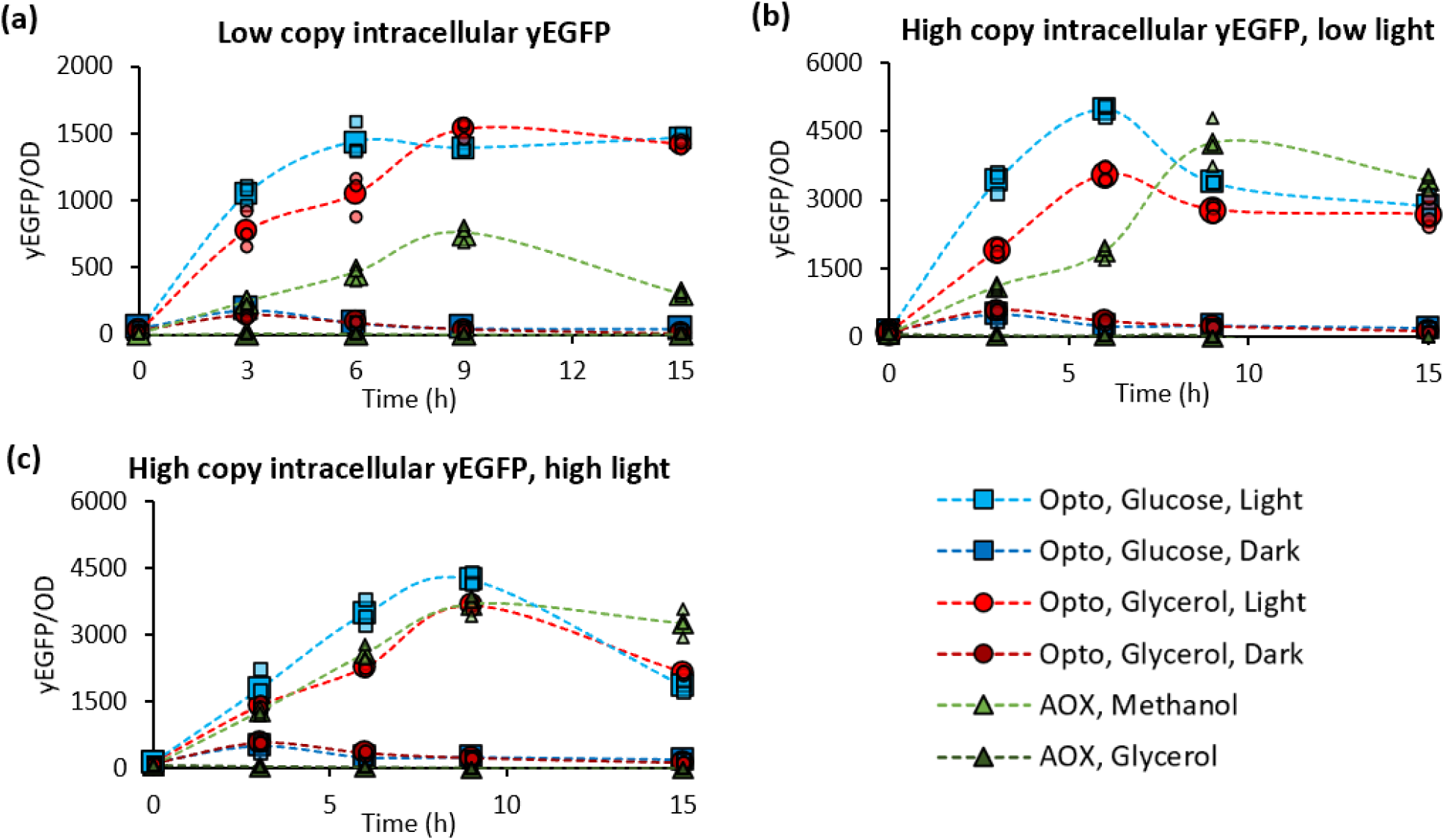
Intracellular yEGFP production with the coupled optogenetic system. Light-induced production of yEGFP in 1% glucose (BMD1) and 1% glycerol (BMG1) is compared to methanol induction of P_AOX1_ at a (a) single copy in 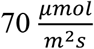 of light as well as at ∼8 copies in (b) 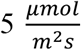 and (c) 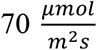 light intensities. Single copy strains are ySMH3-100-3 and ySMH5-100-5. Multicopy strains are ySMH3-1000-3 and ySMH5-1000-3. Large icons represent the mean values, while the smaller points show each of the three independent biological replicates. Dashed lines connect the mean values to aid visualization of trends.

Twelve random colonies from each of the optogenetic and methanol-induced strains were analyzed by qPCR, and strains with single copies (ySMH3-100-3 and ySMH5-100-5) and approximately 8 copies (ySMH3-1000-3 and ySMH5-1000-3) were chosen for further analysis (Supplementary Fig. S1).

To compare expression between the strains, we first inoculated them into 1 mL of buffered minimal medium in a 24-well plate (USA Scientific CC7672-7524) and grew them in the dark for 20 h at 30 °C with shaking at 200 rpm (New Brunswick Innova 2300 Platform Shaker). This growth was performed in buffered minimal glycerol medium BMG1 for the P_AOX1_-driven ySMH5 strains, and in both BMG1 and buffered minimal dextrose medium BMD1 for the ySMH3 optogenetic strains (BMG1: 100 mM potassium phosphate, pH 6; 1.34% yeast nitrogen base; 0.4 μg/mL biotin; 1% glycerol. BMD1: 100 mM potassium phosphate, pH 6; 1.34% yeast nitrogen base; 0.4 μg/mL biotin; 1% glucose). After the overnight growth, the plates were centrifuged at 234*g* (Sorvall Legend XTR Centrifuge) for 5 min and the cultures were resuspended in buffered minimal medium with no carbon source to remove any residual glucose or glycerol (BMX: 100 mM potassium phosphate, pH 6; 1.34% yeast nitrogen base; 0.4 μg/mL biotin). We measured the OD_600_ of these cultures in a TECAN plate reader (Infinite F Plex) and diluted the cells into three identical 48-well plates to an OD_600_ of 1. The optogenetic ySMH3 cells were diluted into both BMD1 and BMG1 media (depending on the carbon source in which they were grown) to test in both 1% glucose and glycerol, while the P_AOX1_-driven ySMH5 cells were diluted into buffered minimal methanol BMM1 or BMG1 media to test protein expression in 1% methanol and glycerol (BMM1: 100 mM potassium phosphate, pH 6; 1.34% yeast nitrogen base; 0.4 μg/mL biotin; 1% v/v methanol). The plates were incubated at 30 °C with 200 rpm shaking (New Brunswick Innova 2300 Platform Shaker), and each plate was exposed to a different light intensity (darkness, 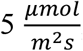, and 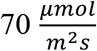). All OD_600_ and yEGFP measurements were obtained using the TECAN plate reader. yEGFP excitation and emission wavelengths of 485 nm and 535 nm were used for fluorescence data, while cell density was measured at a wavelength of 600 nm. Measurements were taken at the indicated times (Fig. 2, Supplementary Fig. S5 and S6), and plates were immediately returned to 30 °C with 200 rpm shaking after each measurement. To avoid unwanted activation of the optogenetic system in the dark plate, this plate was transported in aluminum foil for each timepoint to minimize exposure to ambient light. Unlike the subsequent experiments in shake flasks or the bioreactor, these tests were performed as batch experiments with no further additions of the carbon source after the time of induction.

For the pulsing experiment (Supplementary Fig. S7), ySMH3-100-3 was inoculated into both BMD1 and BMG1 medium in a 24-well plate and grown in the dark for 20 h at 30 °C with shaking at 200 rpm. Cells were then centrifuged at 234*g* as mentioned above for 5 min and resuspended in fresh BMD1 and BMG1 medium at an OD_600_ of 0.1 in four identical 24-well plates. Each plate was subjected to a different light schedule (darkness, 10s ON/100s OFF, 1s ON/100s OFF and constant light) with all panels illuminating at an intensity of ∼70 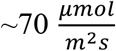. The pulsed light schedules were regulated using a Nearpow Multifunctional Infinite Loop Programmable Plug-in Digital Timer Switch (Model number CECOMINOD036912). OD_600_ and yEGFP were measured after 8 h in each light condition at 30 °C and 200 rpm shaking. Measurements for this experiment were obtained with an Infinite M200PRO TECAN plate reader.

When converting the raw data from these experiments into a yEGFP/OD_600_ or total fluorescence value, we accounted for autofluorescence of the media and cells as well as light-bleaching effects. For yEGFP/OD_600_ values, we used the formula below which has previously been reported for quantifying fluorescence with a similar plate reader^29^ (Equation 1). The wild-type NRRL Y-11430 strain was used as the ‘no yEGFP’ negative control and was subjected to the same conditions (abbreviated as ‘cond’) as the other strains.

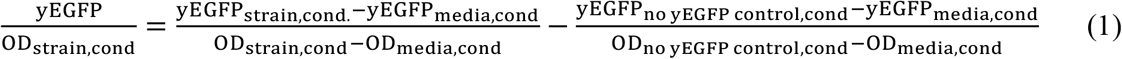

To quantify total yEGFP production, the autofluorescence of the NRRL Y-11430 control was simply subtracted from the measured value of the strain being measured, as in Equation 2.

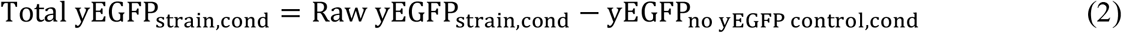

### Testing effect of EL222 copy number on growth and production

To examine how EL222 copy number impacts the behavior of optogenetic strains (Fig. 3, Supplementary Fig. S8 and S9), we first created a parent strain, ySMH115-14, which contains only the P_C120_-yEGFP portion of the optogenetic system (SMH6). To mimic the high copy strains that are generally used in *K. phaffii* bioprocesses, we selected this strain from a plate containing 1000 μg/mL zeocin. We then transformed this strain with a plasmid containing P_ADH2_-EL222 and a G418 marker (SMH139), and plated on 100, 500, and 1000 μg/mL G418. Several colonies were selected from these plates and analyzed with qPCR to identify a set of strains with a wide range of EL222 copy numbers (ySMH193) (Supplementary Fig. S2). This analysis led to the selection of ySMH193-11, 6, and 9 which contain 1, 3, and 8 copies of EL222, respectively.

**Figure 3.**
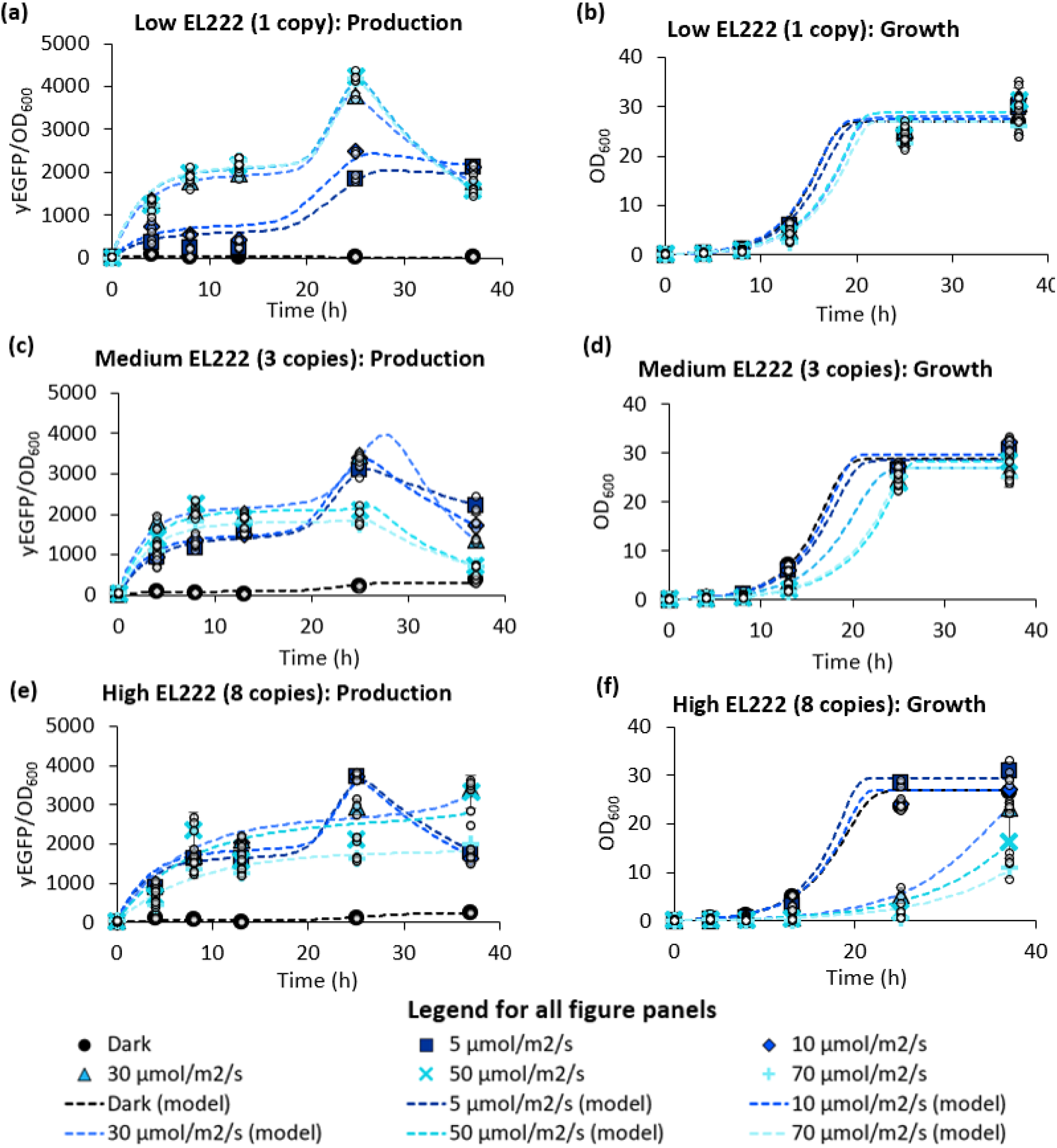
Characterizing the effect of EL222 copy number and light intensity on growth and production. yEGFP-producing strains with different levels of EL222 (ySMH193-11, 6, and 9) were tested at several light intensities to identify a suitable condition for (a, c, e) production and (b, d, f) growth in BMD1 medium. Strains containing (a, b) one, (c, d) three, or (e, f) eight copies of EL222 were examined, with the higher copy strains exhibiting greater sensitivity to light. Mean values are depicted as data points, with error bars indicating the standard deviation from three independent replicates, while the small white dots show each of the three independent biological replicates. Dashed lines illustrate the predicted dynamics generated by the hybrid model with Gaussian-process-predicted parameters. The predicted glucose concentrations are shown in Supplementary Figure S20.

To test the growth and production of these strains, we first inoculated them in BMD1 medium in a 24-well plate, along with the NRRL Y-11430 wild-type and ySMH115-14 no-EL222 controls. This plate was incubated in the dark for 20 h at 30 °C with 200 rpm shaking. The cultures were then centrifuged at 234*g* as above for 5 min and resuspended in fresh BMD1. Cells were diluted into six identical 48-well plates at an OD_600_ of 0.1, as measured by a TECAN plate reader (Infinite F Plex). We exposed each plate to a different light intensity (darkness, 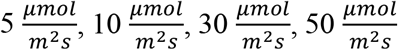 or 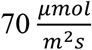) with yEGFP and OD_600_ measurements taken at the indicated timepoints (Fig. 3, Supplementary Fig. S8). For timepoints where the OD_600_ exceeded 8, we diluted a sample of the cells into a separate 48-well plate to measure within the linear range of the instrument. As with the other tests for intracellular yEGFP production, yEGFP/OD_600_ and total yEGFP were quantified using Equations 1 and 2.

### Gaussian-process-supported modeling of yEGFP production

To model the dynamics of protein production under varying EL222 copy number and light intensity, we considered four differential equations, as stated below. As proof of concept, we focused on yEGFP production, although the model can be adapted to other types of recombinant proteins. The model follows:

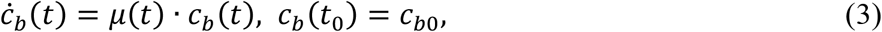

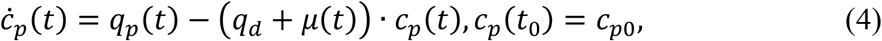

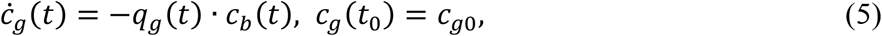

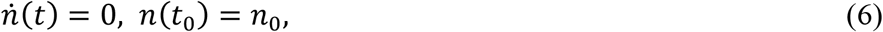

where *c*_*b*_ ∈ ℝ is the cell density (in arbitrary units of OD), *c*_*p*_ ∈ ℝ is the intracellular protein concentration (in arbitrary units of yEGFP), *c*_*g*_ ∈ ℝ is the glucose concentration (in g/L), and *n* ∈ ℕ is the EL222 copy number. Despite being denoted as a dynamic state, the EL222 copy number is a natural number and remains constant throughout the process, as it depends only on the initial condition (i.e., the selected strain). The process time is represented by *t* and the initial process time by *t*_0_. Henceforth, we will omit the time dependency of the variables.

Here, *µ, q*_*p*_, *q*_*d*_, and *q*_*g*_ are appropriate macro-kinetic kinetic functions:

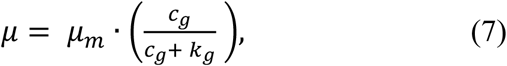

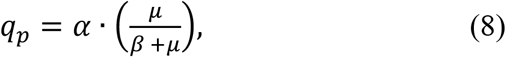

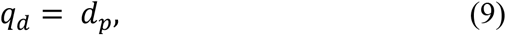

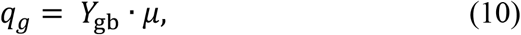

with parameters *µ*_*m*_, *k*_*g*_, *α, β, d*_*p*_, and *Y*_gb_. The specific growth rate, *µ*, follows a hyperbolic function of *c*_*g*_; the intracellular protein production rate, *q*_*p*_, follows a hyperbolic function of *µ*; the intracellular protein degradation rate, *q*_*d*_, is a constant; and the specific glycerol uptake rate, *q*_*g*_, is linearly coupled to *µ*. The model parameters, collected in 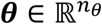, are a function of the light intensity *I* ∈ ℝ and the EL222 copy number. Therefore, 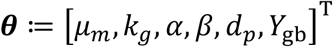 and ***θ*** *= f*_*θ*_ (*I, n*), where 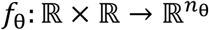 is a vector-valued function mapping light intensity and EL222 copy number to ***θ***. Each model parameter *θ*_*i*_ ∈ ***θ*** is modeled as the mean *function* of a multi-input single-output Gaussian process^45^ 𝒢 𝒫_𝒾_, thereby linking light intensity and EL222 copy number to the dynamic behavior of the system.

Let us denote 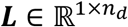 and 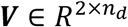 as the training datasets for the output label (*θ*_*i*_) and input features (*I* and *n*) of 𝒢 𝒫_𝒾_, respectively. To populate these training data sets, we conducted a parameter estimation routine for all batch experiments, each corresponding to a specific EL222 copy number and light intensity condition. The parameter estimation was performed using the particle swarm algorithm in COPASI^77^. We fitted the model using the experimental data of the initial glucose concentration and the dynamic profiles of OD_600_ and protein production under different conditions of light intensity and copy numbers.

The training of the Gaussian processes and the numerical integration of the hybrid model, with embedded Gaussian-process-predicted parameters, were performed using the HILO-MPC Python’s library^78^. Overall, 𝒢 𝒫_𝒾_ follows a normal distribution 𝒢 𝒫_𝒾_ (***v***) ∼ 𝒩 (*m*(***v***), *κ*(***v, v***)) with a mean function 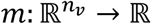 and a kernel (covariance) function 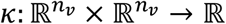. The vector ***v*** ≔ [*I, n*]^T^ ∈ ℝ^2^ represents the features of 𝒢 𝒫_𝒾_. The Gaussian process models a *true function* under normally distributed measurement noise 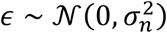, with zero mean and noise variance 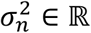. For *n*_*d*_ data entries, the Kernel matrix 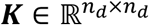 captures the neighborhood of data points in the feature space. Each element (*i, j*) of the Kernel matrix, given two inputs ***v***_*i*_ ∈ ℝ^2^ and ***v***_*j*_ ∈ ℝ^2^, corresponds to the kernel function *κ*(***v***_***i***_, ***v***_***j***_). In this work, we used the Matérn–5/2 kernel function^45^ 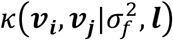, with signal variance 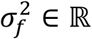 and length-scale ***l*** ∈ ℝ^2^ (given automatic relevance determination for the two features). The hyperparameters of the Gaussian process, 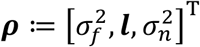, were determined by maximizing the log marginal likelihood 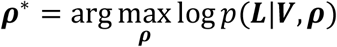. The conditional *posterior* probability for a test input ***v***^*^, considering a Gaussian process with optimal hyperparameters, is normally distributed *p*(𝒢 𝒫_𝒾_ (***v***^*^)|***V, L***) ∼ 𝒩 (*θ*_*i*_, ϒ_*i*_), where the predicted mean is the value of a model parameter *θ*_*i*_, and ϒ_*i*_ ∈ ℝ represents the variance of the prediction. For simplicity of training, the Gaussian processes in this work were trained using the HILO-MPC’s default *prior* mean of zero. However, the prior mean can be reconsidered to enhance model generalizability on unseen data.

### Co-expressing ROS scavengers with EL222

We tested co-expression of EL222 with a total of three ROS scavengers native to *K. phaffii* (*CTA1* catalase, *SOD1* superoxide dismutase, and *PMP20* peroxisomal membrane protein). The sequences for all three genes were identified in previous studies^48,79^ and amplified directly from the *K. phaffii* genome when cloning. In the case of *CTA1*, the last five amino acids were trimmed to remove the putative peroxisomal targeting tag and permit cytosolic expression.

To construct strains co-expressing these scavenger enzymes with EL222, we first knocked out *HIS4* from the wild-type NRRL Y-11430 strain using CRISPR-Cas9 (SMH60) to open a third selection marker, beyond zeocin and G418 resistance (ySMH68). This strain was then transformed with a cassette expressing yEGFP under P_AOX1_ (SMH158, linearized with PmeI to target the AOX1 locus) to form ySMH196 (plated on 100 μg/mL zeocin), and subsequently transformed with a construct containing P_ADH2_-EL222 (SMH139, linearized with ApaI to target the *ADH2* locus) to form ySMH197 (plated on 1000 μg/mL G418). This ySMH197 strain served as the parent strain for ROS scavenger integration, targeting the *HIS4* locus by linearizing SMH219, SMH229, and SMH230 with KasI and yielding ySMH207 (*CTA1*), ySMH209 (*SOD1*), and ySMH210 (*PMP20*), respectively. These ROS scavengers were constitutively expressed from the P_GAP_ promoter.

To test whether co-expression of the scavengers could offset the phototoxic effects of EL222, we inoculated four colonies of ySMH207, ySMH209, and ySMH210 into 500 μL BMD1 medium supplemented with 0.3 mM L-histidine in a 48-well plate, along with ySMH196 (no-EL222 control) and ySMH197 (no scavenger control). Because the ySMH196 and ySMH197 control strains are auxotrophic for L-histidine, this supplemented media was used for all wells to keep conditions consistent between strains (despite ySMH207, ySMH209, and ySMH210 being prototrophic for L-histidine). This plate was grown overnight for 20 h in the dark at 30 °C with shaking at 200 rpm. The next day, the plate was centrifuged at 234*g* for 5 min and resuspended in fresh BMD1 medium supplemented with 0.3 mM L-histidine. The strains were diluted in two identical 48-well plates to an OD_600_ of 0.1 in BMD1 supplemented with 0.3 mM L-histidine as measured by a TECAN plate reader (Infinite F Plex). As previously, the L-histidine was added to all cultures to keep media conditions consistent between the prototrophic scavenger-expressing strains and the auxotrophic control strains. Finally, the two plates were incubated at 30 °C with 200 rpm shaking in their respective light conditions (darkness or 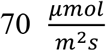 light). Plates were briefly removed at the indicated timepoints (Supplementary Fig. S10) to measure the OD_600_, with samples being diluted in a separate 48-well plate for timepoints where the OD_600_ was above 8 and exceeded the linear range of the instrument. Data is shown from one representative colony of the four tested.

### Intracellular production of Mfp5

Mfp5 from *Mytilus californianus* was codon-optimized for expression in *K. phaffii* (Twist Bioscience) using a previously reported amino acid sequence^49^. We created constructs expressing the gene from both P_AOX1_ and P_C120_ (SMH207 and SMH244). For expression from P_AOX1_, SMH207 was transformed into NRRL Y-11430 cells to yield ySMH186, which was plated on 500 and 1000 μg/mL zeocin agar plates. For optogenetic production, we first created a strain containing a single copy of P_ADH2_-EL222 by transforming NRRL Y-11430 with the linearized SMH139 plasmid (ySMH203-16) (Supplementary Fig. S3). To incorporate the Mfp5 expression cassette, we transformed this strain with SMH244, yielding ySMH238. This transformation was plated on 500 and 1000 μg/mL zeocin agar plates.

To screen the transformants for strong producers, 14 colonies of each strain were inoculated into minimal media (BMG1 for the ySMH186 P_AOX1_-driven strains and BMD1 for the optogenetic ySMH238 strains) in a 24-well plate and grown for 24 h at 30 °C with 200 rpm shaking. This growth was performed in the dark for the optogenetic strain to avoid premature induction. Subsequently, the cultures were centrifuged for 5 min at 234*g* and resuspended in fresh minimal media (BMM1 for ySMH186 and BMD1 for ySMH238). The cell densities were measured using a TECAN plate reader (Infinite F Plex) and equalized to an OD_600_ of 10 in their respective media in 24-well plates. These plates were incubated for 24 h at 30 °C with shaking at 200 rpm, with the optogenetic plate exposed to blue light at an intensity of 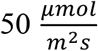. After this period, cells were resupplied with their respective carbon sources to a final concentration of 1% (using 100% methanol for ySMH186 or 40% glucose for ySMH238) and returned to 30 °C for an additional production period of 24 h. We then lysed the cultures at an equal cell mass using a standard trichloroacetic acid (TCA) lysis protocol^80^ and performed SDS-PAGE followed by western blot (see details below) to identify the best performing colonies of each strain (Supplementary Fig. S11).

After identifying the highest producing strains (ySMH186-9 and ySMH238-9), production from these strains was more rigorously compared in triplicate in Erlenmeyer flasks. Each strain was inoculated into 1 mL of minimal media (BMG1 for ySMH186-9 and BMD1 for ySMH238-9) in a 24-well plate and grown for 20 h at 30 °C with 200 rpm shaking. After this growth, 400 μL of each culture was inoculated into 40 mL of fresh media (BMG1 for ySMH186-9 and both BMG1 and BMD1 for ySMH238-9) in 500 mL Erlenmeyer flasks for a second overnight growth. As previously, we grew the optogenetic cells in the dark. We then centrifuged the cells at 234*g* for 5 min and resuspended in 40 mL fresh minimal media (BMM1 for ySMH186-9 and both BMG1 and BMD1 for ySMH238-9). The cell densities were measured using an Eppendorf spectrophotometer (Eppendorf BioSpectrometer basic) and μCuvette (Eppendorf μCuvette G1.0). To ensure the measured cell densities were within the linear range of the instrument (OD_600_<10), 0.25 mL samples of each culture were first diluted 1:5 in fresh media before measurement. Then, additional fresh media was added to each culture such that they were all at an OD_600_ of 10 (BMM1 for ySMH186-9 and both BMG1 and BMD1 for ySMH238-9). This step ensured that the cell density at the time of induction would be equal for all conditions. After equalizing the cell densities to an OD_600_ of 10, 12.5 mL of each culture was transferred to three identical 125-mL Erlenmeyer flasks (nine flasks total), which were then incubated at 30 °C with shaking at 200 rpm. The flasks with optogenetic strains were exposed to light at an intensity of 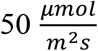 using the flask illuminator platform (Supplementary Fig. S4), while the P_AOX1_-driven strain was exposed to ambient light. After 24 h of incubation, the cells were resupplied with their respective carbon sources to a concentration of 1% (using 100% methanol, 40% glucose, or 60% glycerol) and returned to 30 °C for an additional period of 24 h. We lysed equal cell masses of each culture using the TCA protocol^80^ and quantified production using a western blot and ImageJ analysis (see details below) (Fig. 4). To confirm equal loading of samples and avoid bias in the results, we stained the membrane using Coomassie blue R-250 as previously described^81^ (Supplementary Fig. S12).

**Figure 4.**
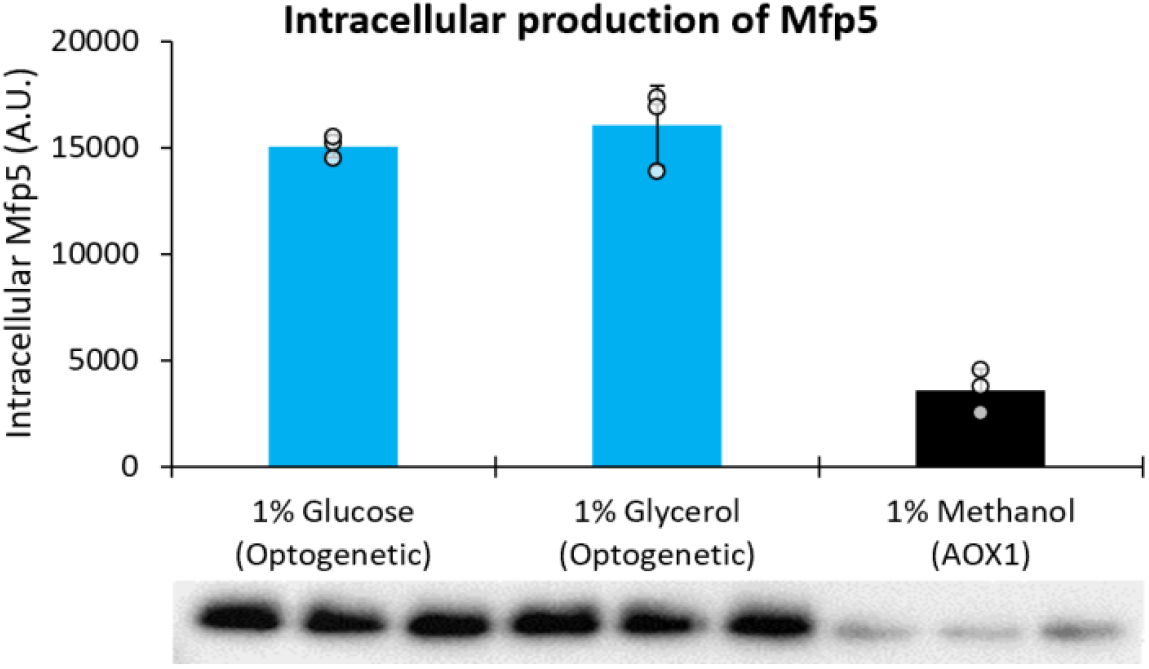
Intracellular production of *M. californianus* Mfp5 with optogenetics and P_AOX1_. Production of Mfp5 is compared between P_AOX1_ and the decoupled optogenetic system using 1% glucose or glycerol for the optogenetic strain (ySMH238-9) and 1% methanol for the P_AOX1_ strain (ySMH186-9). Optogenetic cultures were exposed to light at 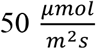 intensity and fermentations were carried out for 48 h. Pixel intensity in the western blot bands was analyzed using ImageJ to generate quantitative values for Mfp5 production. Mean values after 48 h are shown, and error bars represent the standard deviation of three biological replicates.

### Protein quantification through SDS-PAGE and western blot analysis

Production of Mfp5 and the three secreted targets were analyzed through SDS-PAGE followed by western blots. For the intracellular production strains, samples were lysed using a standard TCA protocol^80^ followed by resuspending in a SDS sample buffer^82^. For secreting strains, the final cultures were centrifuged for 5 min at 234*g* and the supernatant was mixed with the sample buffer^82^. We boiled samples for 5 min at 100 °C using a heat block (VWR), loaded onto 12% SDS-PAGE gels, and resolved. We used a loading volume of 15 μL for all SDS-PAGE gels.

After resolving, we analyzed by western blot using established protocols. Proteins were transferred onto a PVDF membrane using a Trans-Blot Turbo Transfer System (Bio-Rad). All proteins analyzed by western blot contained a C-terminal His tag which was assayed using THE™ His Tag Antibody [HRP], mAb (Genscript) with a dilution of 1:10000. Blots were revealed using Clarity ECL substrate (Bio-Rad) and imaged with a Bio-Rad ChemiDoc MP Imaging System with Image Lab software, using the chemiluminescence protocol. After imaging, blots could be quantified by using Fiji ImageJ software^83^ to correlate pixel intensity to protein production.

### Verifying phototoxic effect of EL222 in secretion strains

We tested whether phototoxicity would affect secreted production by co-expressing EL222 in a strain that secretes yEGFP from P_AOX1_ using the α-factor secretion tag (Fig. 5). To create this strain, we first transformed NRRL Y-11430 with the plasmid for yEGFP secretion (SMH158) and plated on a 1000 μg/mL zeocin agar plate, yielding ySMH154-1. This strain was then transformed with a plasmid containing EL222 (SMH139) and plated on 1000 μg/mL G418 to create ySMH185. After creating these two strains, we inoculated each into 1 mL BMG1 medium in a 24-well plate and incubated for 20 h in the dark at 30 °C with 200 rpm shaking. After this period, 150 μL of each culture was inoculated into 15 mL of BMG1 in 125 mL Erlenmeyer flasks and grown again for 20 h at 30 °C and 200 rpm shaking in the dark. The cultures were then centrifuged for 5 min at 234*g* and resuspended into 15 mL BMM1 medium. An Eppendorf spectrophotometer (BioSpectrometer basic) was used to measure the cell densities, and the cultures were equalized to an OD_600_ of 10 in BMM1 and pipetted in triplicate to five identical 24-well plates. Each plate was incubated 24 h at 30 °C and 200 rpm shaking, with each plate exposed to a different light intensity (darkness, 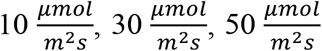, or 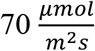). After this period, the cultures were resupplied with methanol to a final concentration of 1% and returned to their respective light conditions for 24 h. Finally, the plates were centrifuged for 5 min at 234*g* and analyzed by dot blot using 350 μL of the supernatant (Fig. 5a). This dot blot was subsequently quantified using Fiji ImageJ software^83^ to convert pixel intensity to relative production (Fig. 5b).

**Figure 5.**
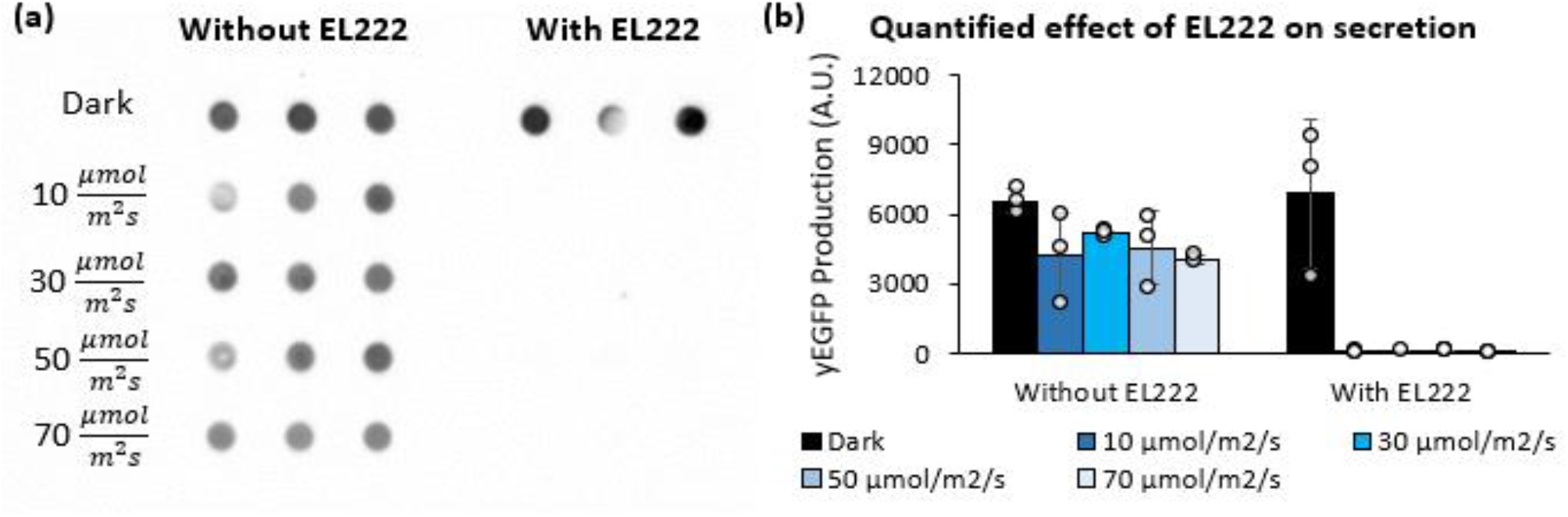
Phototoxic stress of EL222 can inhibit secretion even at low light levels. The phototoxic effect on secretion was verified by co-expressing EL222 in a strain secreting yEGFP from P_AOX1_. (a) Secretion by a non-optogenetic P_AOX1_-driven strain without EL222 (ySMH154-1) and with EL222 (ySMH185-1) at a range of light intensities is shown by dot blot, in which each dot represents a biological replicate induced for 48 h in BMM1 medium. (b) This effect was quantified using ImageJ analysis of the dot blot, in which mean values are shown and error bars represent the standard deviation of the three replicates.

### Dot blot analysis of secreted proteins

Dot blots were used to analyze secretion experiments wherein the number of samples exceeded what could be run on an SDS-PAGE gel (Fig. 5a, Supplementary Fig. S13, S14, and S15). We centrifuged the culture plates for 5 min at 234*g* and filtered 350 μL of supernatant from each well through a nitrocellulose membrane in a Bio-Dot Microfiltration Apparatus (Bio-Rad) using the recommended protocol for the equipment. After filtration, membranes were incubated for 1 h at room temperature in a blocking buffer (1X TBS, 0.1% Tween-20 with 5% w/v nonfat dry milk) and assayed using THE™ His Tag Antibody [HRP], mAb (Genscript) at a 1:10000 dilution. After incubating with the antibody for 1 h at room temperature, we washed the membranes four times for 10 min with TBST (1X TBS, 0.1% Tween-20). Finally, we imaged the blots with Clarity ECL substrate (Bio-Rad) on a Bio-Rad ChemiDoc MP Imaging System using the chemiluminescence protocol.

### Secretion of yEGFP, SR18, and β-lactoglobulin using optogenetics and P_AOX1_

Secreted production of three proteins was compared between P_AOX1_ and two versions of the decoupled optogenetic system (differing by EL222 copy number). The methanol-induced strains were created by transforming NRRL Y-11430 with a construct secreting the gene of interest from P_AOX1_ using the α-factor secretion tag, yielding ySMH154 (yEGFP), ySMH223 (SR18), and ySMH51 (β-lactoglobulin), all using PmeI to linearize the plasmids. We constructed the optogenetic strains by transforming ySMH203-16 (single copy of EL222) and ySMH203-2 (five copies of EL222) with plasmids linearized with PmeI to secrete the gene of interest from P_C120_ using the same α-factor secretion tag. The resulting strains with one copy of EL222 were ySMH217 (yEGFP), ySMH230 (SR18), and ySMH224 (β-lactoglobulin), while the strains with five copies of EL222 were ySMH219 (yEGFP), ySMH231 (SR18), and ySMH226 (β-lactoglobulin). All transformations were plated on 500 and 1000 μg/mL of zeocin. We screened 94 colonies of each strain in a 96-well format in a deep-well plate (USA Scientific). For the methanol-induced strains, each colony was inoculated into 500 μL of BMG1 medium in the deep-well plate, with the last two wells saved for wild-type and blank negative controls. After one day incubation period at 30 °C and shaking at 300 rpm, the plates were centrifuged at 234*g* for 5 min and the supernatant was replaced with 500 μL of fresh BMM1 medium. The cells were induced for 48 h at 30 °C with 300 rpm shaking, with methanol added after 24 h to return the final concentration to 1% (assuming all the initial methanol had been consumed in the first 24 h). After the production phase, the plates were again centrifuged for 5 min at 234*g* and 350 μL of supernatant from each well was analyzed by dot blot (Supplementary Fig. S13a, S14a, and S15a). The highest-producing colony of each strain was selected for future experiments at the flask-scale.

To screen the optogenetic strains, 94 colonies of each strain were inoculated into 500 μL BMD1 medium in the deep-well plates and grown in the dark for one day at 30 °C with 300 rpm shaking. The plates were then centrifuged for 5 min at 234*g* and resuspended in fresh BMD1 medium. We placed the plates in their respective suitable light conditions (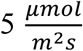 for the 5-copy EL222 strains and 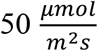 for the 1-copy EL222 strains) to produce for 48 h at 30 °C with shaking at 300 rpm. Glucose was added to a final concentration of 1% at the 24-h point of this incubation (assuming all the initial glucose had been consumed in the first 24 h). Finally, we centrifuged the plates at 234*g* for five min and analyzed by dot blots as with the methanol-induced strains (Supplementary Fig. S13b-c, S14b-c, and S15b-c).

With the best producer of each strain identified, we then performed tests in shake flasks to compare production between optogenetics and P_AOX1_ in triplicate. First, each strain was inoculated into 1 mL of minimal media (BMG1 for ySMH154-C5, ySMH223-G5, and ySMH51-D2 and BMD1 for ySMH217-A2, ySMH219-D7, ySMH230-A6, ySMH231-G2, ySMH224-A11, and ySMH226-C1) in a 24 well plate and grown for 20 h at 30 °C with 200 rpm shaking in the dark. After this growth, 400 μL of each culture was inoculated into 40 mL of fresh media (BMG1 for the methanol-induced strains and both BMG1 and BMD1 for the optogenetic strains) in 500 mL Erlenmeyer flasks for a second overnight growth in the dark. After 20 h, we centrifuged the cultures for 5 min at 234*g* and resuspended in 40 mL fresh media again (BMM1 for the methanol-induced strains and both BMG1 and BMD1 for the optogenetic strains). We measured the cell densities with an Eppendorf spectrophotometer (BioSpectrometer basic) and equalized to an OD_600_ of 10 in their respective media. 12.5 mL of each culture was transferred to three identical 125-mL Erlenmeyer flasks, which were then incubated at 30 °C with shaking at 200 rpm. The optogenetic flasks were exposed to light at an intensity of 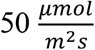 for the 1-copy EL222 strains and 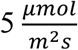 for the 5-copy EL222 strains using the illuminator platform (Supplementary Fig. S4). After 24 h of incubation, the cells were resupplied with their carbon sources to a concentration of 1% and returned to 30 °C for an additional 24 h. We harvested the fermentations by centrifuging the cultures for 5 min at 234*g* and analyzing the supernatant using quantitative western blots (Fig. 6, Supplementary Fig. S16, S17, and S18).

**Figure 6.**
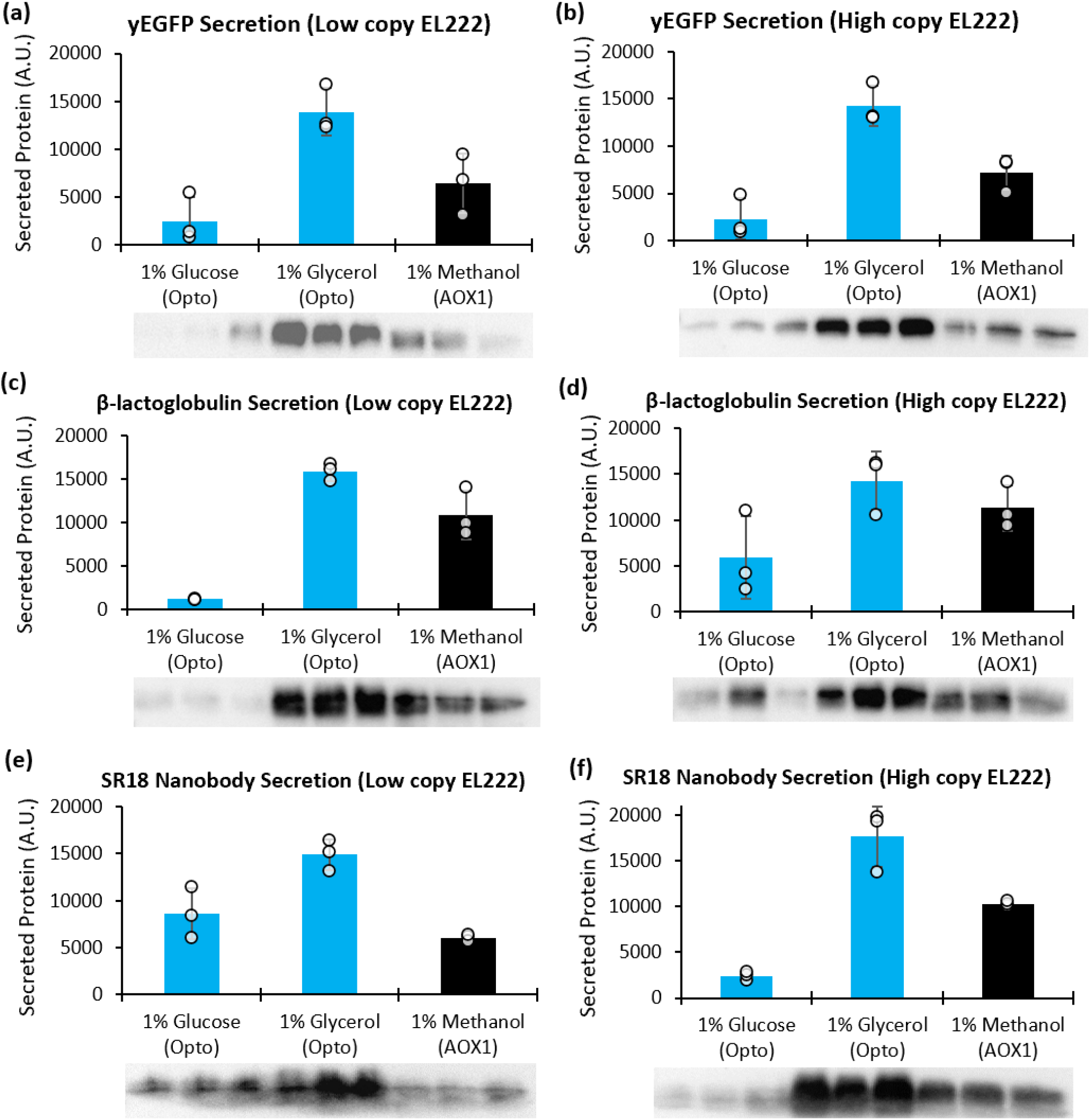
Optogenetic secretion of diverse proteins compared to P_AOX1_. Secretion of (a, b) yEGFP, (c, d) β-lactoglobulin dairy protein, and (e, f) the SR18 nanobody was tested in triplicate with the best optogenetic and methanol-induced strains. For the optogenetic strains, we tested the best producers from (a, c, e) a low copy EL222 background at 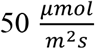 light intensity and (b, d, f) a high copy EL222 background at 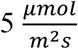 intensity in both BMD1 and BMG1 media. P_AOX1_-driven strains were induced in BMM1 medium. Western blots were quantified using ImageJ, and bar graphs depict mean values with error bars representing the standard deviation of three replicates.

### Scale-up of production in a 2-L bioreactor

Production of the SR18 nanobody was scaled to the 2-L bioreactor level using the ySMH223-G5 (methanol-induced) and ySMH230-A6 strains (optogenetic with 1 copy of EL222), which were identified as the best producers by dot blot screens. Both strains were first inoculated into 5 mL of YPD media with 100 μg/mL zeocin and grown in the dark for 20 h with 200 rpm shaking. We used 2 mL of the grown cultures to inoculate 50 mL of YPD in a 500 mL Erlenmeyer flask containing 100 μg/mL zeocin and grew them again for 20 h in the dark. After this growth, the cells were concentrated by centrifuging for 5 min at 234*g* and removing 25 mL of the supernatant, after which the OD_600_ was measured using an Eppendorf spectrophotometer (BioSpectrometer basic). To ensure the measured cell densities were within the linear range of the instrument (OD_600_<10), a 0.1 mL sample of each culture was first diluted 1:10 in fresh media before measurement. The bioreactor experiments followed a pre-defined recipe, arbitrarily chosen and not derived from any design of experiments or optimization procedure. However, it serves the purpose of providing preliminary insights into the process dynamics at this scale.

We set up a BioFlo120 system with a 2-L bioreactor (Eppendorf, B120110001) and added 1 L of basal salts medium (BSM) supplemented with 4.35 mL PTM1 trace salts^84^ and 4% w/v glycerol, along with 100 μL of Antifoam 204 (Sigma-Aldrich). The reactor was set to 30 °C with a pH of 5.0, which was maintained using 14% ammonium hydroxide. After reaching these setpoints, the dissolved oxygen probe was calibrated and set to a minimum percentage of 30% by adjusting the agitation speed (200-600 rpm) and air flow rate (0.1-3 SPLM). Cell density was measured using a noninvasive biomass sensor with a wide linear range from less than 0.1 to greater than 300 OD units (bug lab BE2100). We then inoculated the reactor with the concentrated culture (either ySMH223-G5 or ySMH230-A6) to an OD_600_ of 1. The cells were grown for ∼24 h in the repressive conditions (darkness in the case of the optogenetic ySMH230-A6 strain) until a spike in the dissolved oxygen was observed, which signaled the end of the batch phase. After the batch phase, a brief semi-batch phase was performed (still in repressive conditions for both the optogenetic and non-optogenetic bioreactor experiments) by feeding 50% w/v glycerol supplemented with 12 mL/L of PTM1 for 4 h at a rate of 6 mL/h. A spike in the dissolved oxygen was observed after stopping the feed, indicating full consumption of glycerol.

To induce production in the P_AOX1_-controlled strain, 100% methanol supplemented with 12 mL/L PTM1 was fed to the reactor at an initial rate of 3.6 mL/h. After 4 h at this rate, the feed was turned off until another spike in the dissolved oxygen was observed, after which the flow rate was increased to 7.2 mL/h. The methanol was fed at this higher flow rate for 48 h (the remainder of the fermentation). To prevent excess foaming, 100 μL of Antifoam 204 was added approximately every 12 h after the onset of induction. Samples of 5 mL were taken at each timepoint post-induction for analysis using Coomassie staining of SDS-PAGE gels and Bradford assays as previously described^85,86^.

To induce production in the optogenetic bioreactor runs, an LED panel positioned under the reactor was illuminated, and the glycerol mixture (50% w/v glycerol supplemented with 12 mL/L PTM1) was fed at a rate of 4.5 mL/h. This feed rate was selected to provide an equivalent amount of carbon source on a mass basis as the methanol-induced experiments. After 4 h, the flow rate was increased to 9 mL/h for the remainder of the fermentation. Antifoam 204 was added, and samples were taken as aforementioned in the methanol-induced bioreactor runs. Samples were analyzed using Coomassie staining of SDS-PAGE gels and Bradford assays as previously described at the same time and with the same reagents as the methanol-induced cultures^85,86^. For the Bradford assays, bovine serum albumin (BSA) (Sigma-Aldrich) was used to generate the standard curve.

## Results

To establish optogenetic control of gene expression in *K. phaffii*, we constitutively expressed EL222 using the native P_ADH2_ promoter with the gene of interest downstream of the P_C120_ promoter^41^. We selected the P_ADH2_ promoter to express EL222 due to its previously reported moderate constitutive behavior in both glucose and glycerol^42^. We describe two light-activated system designs that allow for either coupled or decoupled integration of EL222 and the gene of interest (Fig. 1a). The coupled system, which benefits from simplicity, employs a single vector with a zeocin resistance selection marker, and thus integrates an equal number of copies of EL222 and the gene of interest. The decoupled system integrates these genes separately, which allows for independently selected copy numbers of EL222 and the gene of interest using G418 and zeocin, respectively. Both systems offer different advantages in the development of strains for two-phase processes characterized by a dark growth phase and light-induced production phase (Fig. 1b).

### Development and characterization of an optogenetic inducible system in *K. phaffii*

Under the tested conditions, the simpler coupled system (Fig. 1a) achieves stronger induction of intracellular protein production than the methanol-induced P_AOX1_ promoter. Comparing the normalized expression of yEGFP 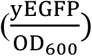 in strains containing a single copy of the expression cassette, using the coupled optogenetic system or P_AOX1_ (Supplementary Fig. S1) shows that blue light induction of EL222 yields approximately twice as much yEGFP than methanol induction (Fig. 2a). For this test, we used an intensity of 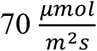, which falls within the typical range for microbial optogenetic experiments^29,31^. In this case, light induction is also faster than methanol induction and shows similar strength in both glucose and glycerol. Because cells grow faster and to a higher density in these carbon sources than in methanol^43,44^, light induction also leads to substantially higher overall intracellular protein production levels (Supplementary Fig. S5a). Total production from the optogenetic strain continues to increase at 15 h, whereas P_AOX1_ plateaus by that time, suggesting the coupled system could produce even more protein in longer cultivations (Supplementary Fig. S5a). Additionally, optogenetic induction is tunable to a wide range of expression levels by adjusting the light dosage (Supplementary Fig. S7), offering flexibility for intermediate expression strengths. Having found it to be effective at a single integrated copy, we next explored how this system performs at the higher copy numbers, more typical of production processes.

When integrated at a higher copy number, the coupled optogenetic system remains competitive with P_AOX1_ induction levels. Hypothesizing that the higher levels of EL222 expression may increase the light response sensitivity, we identified strains containing approximately 8 copies of either the coupled optogenetic system or P_AOX1_ (Supplementary Fig. S1), and tested expression at both a low 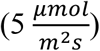 and high light intensity 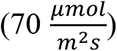. In both conditions, the expression strength when normalized by cell density 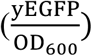 is comparable to P_AOX1_, but light induction is substantially faster (Fig. 2b-c). Interestingly, we found that the optogenetic strain containing a high-copy number of EL222 (and yEGFP) is more productive at the lower light intensity (Fig. 2b-c and Supplementary Fig. S5b-c), suggesting possible phototoxicity at 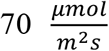 consistent with reduced cell growth at this light intensity (Supplementary Fig. S6). Finally, though production remains strong compared to P_AOX1_ across copy number and light conditions, P_C120_ is slightly leakier, likely due to the lack of a repression mechanism of EL222. Having confirmed that the high-copy optogenetic strain excels at a low light intensity, we sought to clarify the connection between EL222 copy number and light sensitivity.

To test our hypothesis that high EL222 expression increases photosensitivity, we decoupled the copy number of EL222 and recombinant protein (Fig. 1a, bottom). We used this approach to analyze growth and production at a diverse range of EL222 copy numbers and light intensities. To ensure that EL222 copy number was the only difference between strains, we integrated the P_ADH2_-EL222 cassette into a parent strain already containing 12 copies of P_C120_-yEGFP. Transformants were selected and screened for a range of EL222 copy numbers (1, 3, or 8 copies) using qPCR (Supplementary Fig. S2). We then measured growth and fluorescence of each strain at different light intensities (Fig. 3), compared to a control strain lacking EL222 (Supplementary Fig. S8). As our goal was to characterize the relationship between EL222 copy number and light intensity, methanol-induced strains were not tested in this experiment. Consistent with our earlier observations, we found that the strain with one copy of EL222 expresses better at high light intensities 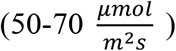, while the strain containing 8 copies is better at the lowest intensities tested, 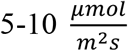 (Fig. 3a and e, Supplementary Fig. S9). Consistent with this trend, an intermediate 3 copies of EL222 reaches its highest activation at a moderate light intensity 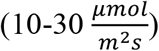 (Fig. 3c, Supplementary Fig. S9). In addition to inhibiting protein production, high light dosages dramatically reduce cell growth in the strain with 8 copies of EL222 (Fig. 3b, d, f). In other words, high light intensities become detrimental only when EL222 is expressed, especially at high levels (Fig. 3 and Supplementary Fig. S8, S10a, and S10b). Regardless of EL222 copy number or light intensity, there is generally a decrease in fluorescence at later times after cell growth plateaus. This is likely due yEGFP gradually being degraded after the carbon source is fully consumed. Together, these observations highlight the importance of selecting the light dosage based on the EL222 expression level and argue against the tenet that higher light doses *always* lead to higher activation. To capture this complex interaction, we developed a hybrid Gaussian-process-supported mathematical model to fit the dynamics of the yEGFP-producing system described above, under varying conditions of EL222 copy number and light intensity. The hybrid model satisfactorily fits the dynamic behavior of cell density and protein concentration. To highlight the data-driven nature of the parameter *functions*, we show in Supplementary Fig. S19 selected parameter values at different light intensities and copy numbers.

We next investigated the phototoxicity of EL222. One hypothesis is that activation of EL222 generates damaging reactive oxygen species (ROS), which could inhibit cell growth and protein production. Thus, higher EL222 levels and light intensities would increase ROS concentrations and exacerbate these effects. Though not previously shown with EL222, other light-responsive proteins containing a LOV domain are known to emit ROS, particularly singlet oxygen, when activated by light^46,47^. To test whether the same is true for EL222, we overexpressed the endogenous superoxide dismutase (*SOD1*), peroxisomal membrane protein 20 (*PMP20*), and catalase (*CTA1*) genes, which code for the main ROS-scavenging enzymes in *K. phaffii*^48^ (Supplementary Fig. S10). Unlike the parent strain housing 7 copies of EL222 without overexpressing an ROS scavenging enzyme (Supplementary Fig. S10b), the strain overexpressing *PMP20* grew equally well with or without light exposure (Supplementary Fig. S10c) while the strains overexpressing *CTA1* or *SOD1* did not improve growth in the light (Supplementary Fig. S10d, e). Given that the main ROS generally emitted by phototoxic LOV domain-containing proteins is singlet oxygen^46,47^, and that Cta1p and Sod1p consume hydrogen peroxide and superoxide respectively, it is not surprising that these enzymes did not protect cell growth from the toxic combination of blue light and EL222. While the ROS substrates for Pmp20p are less characterized than for Sod1p and Cta1p, our results support the hypothesis that ROS emission by EL222 is the cause of its phototoxic effects. This ROS emission, which likely increases with EL222 copy number, offers new insights into improving production by optimizing the levels of EL222 and light. The best strategy to achieve this is to employ the decoupled optogenetic system since it integrates EL222 and the gene of interest independently (Fig. 1a bottom panel). We utilized the decoupled optogenetic system to produce proteins beyond yEGFP, including Mfp5 mussel foot protein, β-lactoglobulin, and a nanobody against SARS-CoV-2.

### Optogenetic induction for intracellular production of Mfp5 mussel foot protein

To demonstrate the capability of optogenetics to control the intracellular production of proteins besides yEGFP, we used the decoupled system to produce the Mfp5 mussel foot protein from *Mytilus californianus*, which has applications in the biomaterials industry^49^. We transformed a strain containing a single copy of EL222 with a gene cassette containing P_C120_-Mfp5 and a zeocin selection marker. To compare light with methanol induction, we also transformed the wild-type strain to express Mfp5 with P_AOX1_ instead of P_C120_ and tested for production with methanol or blue light induction. After screening 14 optogenetic strains induced with blue light (at 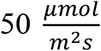 intensity) and 14 methanol-induced strains for Mfp5 production (Supplementary Fig. S11), the top producers of each were compared in triplicate (Fig. 4, Supplementary Fig. S12). Under the tested conditions, we found that Mfp5 production with the optogenetic system is greater than with P_AOX1_.

### Optogenetic induction of secreted proteins

Next, we turned our focus to protein secretion, which often simplifies industrial downstream purification operations. First, we investigated whether the inhibitory effects of high EL222 levels at high light intensities also occur when secreting proteins. To test this, we compared two strains that secrete yEGFP from P_AOX1_, differing only in whether they co-express EL222. In this experiment, EL222 is not actually controlling protein expression, so strains should perform comparably regardless of light intensity if EL222 is not a factor. However, protein secretion from the EL222-expressing strain was severely impaired by any exposure to light (Fig. 5). In contrast, the control strain is relatively unaffected at all intensities tested. These results are consistent with our intracellular experiments (Fig. 3, Supplementary Fig. S10), showing that EL222 activation at suboptimal light conditions affects growth and protein production, whether it is secreted or produced intracellularly. With this insight, we applied the decoupled optogenetic system for protein secretion, carefully selecting light intensities based on the copy number of EL222.

As a first target, we compared yEGFP secretion between optogenetic and methanol-induced strains. For the optogenetic system, we first constructed a set of platform strains with various copy numbers of EL222, which were verified by qPCR (Supplementary Fig. S3). Strains with both low (one) and high (five) copies of EL222 were then transformed with P_C120_-yEGFP and tested for yEGFP secretion at innocuous light intensities: 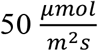 for 1 copy of EL222, and 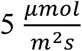 for 5 copies of EL222. For each system, we screened 94 strains by dot blot to identify suitable producers (Supplementary Fig. S13). After selecting the top optogenetic and methanol-induced strains, we compared their yEGFP secretion in triplicate, with the optogenetic strains cultivated (and induced) in media containing glucose or glycerol (Fig. 6a and b, Supplementary Fig. S16). Under the tested conditions, quantitative western blots show that the optogenetic strains in glycerol media produce more yEGFP than the methanol-induced strain by approximately two-fold. However, unlike with intracellular production, optogenetic secretion in glucose is weaker than P_AOX1_. The cause of this discrepancy is not clear but is consistent across the low- and high-EL222 strains. Nevertheless, the strong production in glycerol motivated us to test the secretion of other proteins of higher value.

To show that optogenetic secretion is effective for proteins beyond yEGFP, we used it to produce β-lactoglobulin dairy protein^50^ and the SR18 nanobody which neutralizes the SARS-CoV-2 virus spike protein^51^. As with yEGFP, we used dot blots to screen optogenetic and methanol-induced strains and compared the best producers in triplicate (Supplementary Fig. S14 and S15). For both proteins, under the tested conditions, light induction of P_C120_ achieved higher production in glycerol compared to methanol induction of P_AOX1_, regardless of EL222 copy number (Fig. 6c-f, Supplementary Fig. S17 and S18). As previously observed, secretion in glucose is weaker than in glycerol and generally weaker than P_AOX1_. Together, these results show that light can be a powerful alternative to methanol induction for diverse applications, particularly in glycerol media.

### Scale up of light induction to bioreactor

Having achieved strong light induction of protein production in shake flasks, we next tested our system in a lab-scale bioreactor. As the SR18 nanobody is the highest value product of the three targets examined, we selected it as a representative example for scale-up. Although the strain with five copies of EL222 requires less light than the single copy strain, we moved forward with one copy of EL222, as a proof of concept, because it offers a wider range of effective light dosages (Fig. 3, Supplementary Fig. S9). It also constitutes the strictest option to explore the potential challenge of limited light penetration in (large-scale) bioreactors. We first grew cells in the dark, in a 2-L batch bioreactor using glycerol as a carbon source. In addition to controlling light exposure, we also controlled the pH and monitored dissolved oxygen (DO). When the glycerol was fully consumed, we started a glycerol feed to run a fed-batch process, keeping the reactor in the dark for 4 h before illuminating the bottom of the reactor to induce SR18 production (Supplementary Fig. S21). As a control, we also performed a more traditional methanol-induced fed-batch process with an equivalent carbon source feed rate of methanol on a mass basis, keeping identical pH and DO setpoints. Although neither the optogenetic nor methanol-inducible processes were a priori optimized, and therefore cannot be directly compared to determine superiority, these experiments provide a foundation for comparing the dynamics of both induction systems and grant insight towards future process optimization. The optogenetic and methanol-induced strains were similarly productive for the first 16 h, with both secreting approximately 0.5 g/L of SR18; however, under the tested conditions, the optogenetic strain outperforms the methanol-induced process at every subsequent timepoint, ultimately reaching a final titer of 1.5 ± 0.6 g/L by the end of the fermentation, as measured by Bradford assay (Fig. 7). The optogenetic strain likely benefits from the lower oxygen requirement of glycerol metabolism relative to methanol^52^ along with the lower toxicity of this substrate, which allows for faster consumption, lower levels of oxygen, and potentially greater production levels.

**Figure 7.**
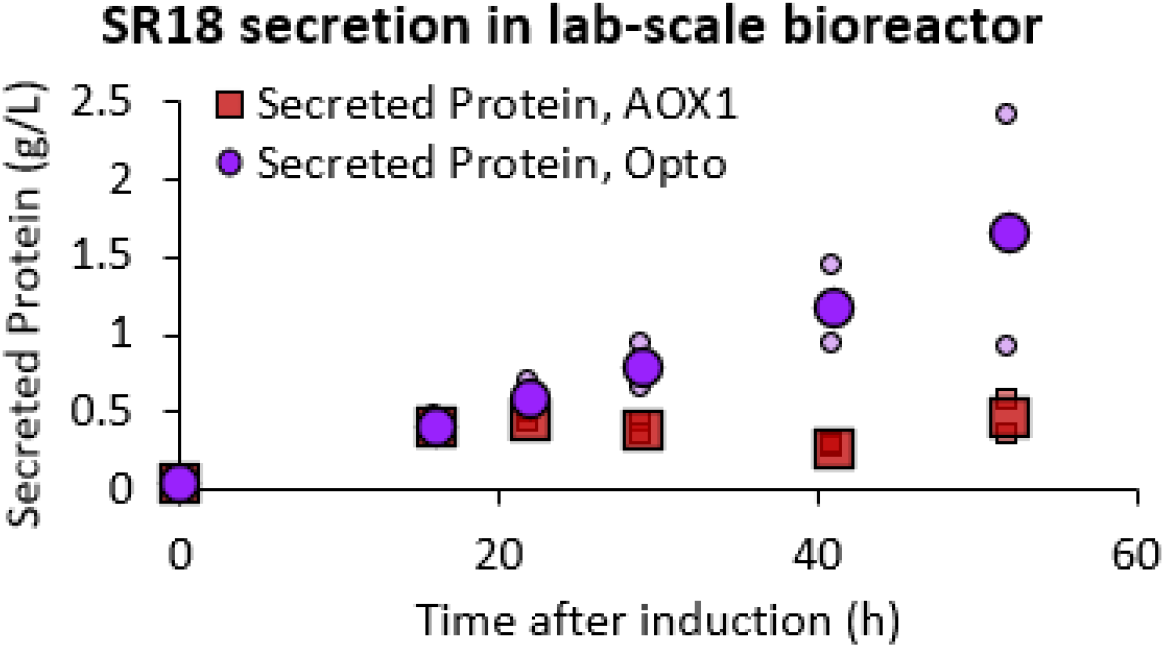
Optogenetic secretion in a bioreactor. The SR18 nanobody was secreted at the bioreactor scale to test optogenetic induction in a larger volume and high cell density. The optogenetic system is compared to a methanol-induced bioprocess in which equal amounts of the carbon sources were fed on a mass basis. Samples taken at the indicated timepoints were analyzed by Bradford analysis to measure the secreted protein titers. Data is shown as mean values, and smaller icons represent individual replicates.

To confirm that most of the secreted protein in these experiments was the intended SR18 nanobody, we paired our Bradford analyses with gel electrophoresis (Fig. 7 and Supplementary Fig. S22). In both systems, as expected, no protein secretion was detected at the onset of induction (during the glycerol phase for P_AOX1_ strain, and in the dark for the optogenetic strain). Yet, this nanobody seems to be unstable given its appearance as multiple bands in the stained gels; a trait to consider if this protein is selected for further bioprocessing. Nevertheless, since this observation occurs in both the optogenetic and P_AOX1_-driven processes, the overall titers remain a fair comparison between the two systems, even if the Bradford analysis may overestimate the absolute quantity of functional protein.

In addition to being two orders of magnitude greater in volume than the shake flask tests, the bioreactor cultures reached exceptionally high cell densities, with the light-induced cells growing more than their methanol-induced counterparts by almost two-fold (Supplementary Fig. S23). Since greater biomass is not necessarily an advantage for secreted proteins due to biomass-product metabolic tradeoff, there is likely an opportunity to further increase the volumetric productivity of the recombinant protein by optimizing the dynamic control strategy to add balance between cell growth and production. Ultimately, the successful production of SR18 by optogenetic induction with only one copy of EL222 (the least light-sensitive strain we developed) shows that light penetration is not an obstacle at least at this scale. Nevertheless, future research should explore the feasibility and performance of the optogenetic approach in larger bioreactor systems, coupled to suitable bioreactor designs to support light-mediated dynamic metabolic control at an industrial scale.

## Discussion

In this study, we report tools and methods for methanol-free induction of recombinant protein production in *K. phaffii* with light. The two general strategies we developed, the coupled or decoupled systems (Fig. 1a), have different advantages: simplicity (coupled) versus versatility in strain design for varying light sensitivities to protein production (decoupled). Both systems are strongly activated in glucose or glycerol, eliminating the need to switch carbon sources mid-process as is the case with the P_AOX1_ promoter. Furthermore, because *K. phaffii* metabolizes glucose and glycerol faster and reaches higher biomass with these carbon sources than methanol, protein production is overall higher and faster using optogenetic induction under the tested conditions (Fig. 2, Supplementary Fig. S5 and S6). Together, the outlined optogenetic methods are promising for inducing protein production and should be considered as potential alternatives to P_AOX1_ or other media-based approaches.

A previous study showed that EL222 is functional in *K. phaffii*^40^, but fell short of exploring conditions in which light induction could be comparable to methanol induction. In contrast, we carefully compared the performance of different optogenetic strain designs with that of P_AOX1_, controlling for critical factors like copy number and optimal light intensity. This type of characterization connecting EL222 expression level to optimal light dosage for gene induction had not been previously reported in yeast or any other organism. However, adjusting light dosage in the context of EL222 copy number proved essential for optogenetic induction of protein production in *K. phaffii* at levels that rival methanol induction. This suggests that similar characterization of other optogenetic systems involving different light-responsive proteins will likely be valuable to expand the capabilities of those systems.

A key finding in our study is the importance of adjusting light intensity depending on the EL222 copy number. Blue light at all intensities we used did not show any phototoxicity to the wild-type strain. However, the higher light doses impair growth and production of both intracellular and secreted recombinant proteins in strains containing high copies of EL222 (Fig. 3 and 5, Supplementary Fig. S9). This observation poses a caveat to the frequently discussed low-toxicity of optogenetics^28,40,53–55^, in that EL222 is only non-toxic below a certain light threshold that depends on EL222 concentration. Our group has shown that maximal light exposure activates EL222 maximally in *S. cerevisiae* without any observable phototoxicity^29– 31,34^, but all strains in those studies contained only one copy of EL222. Consistent with these trends, some studies report poor growth when strongly expressing other optogenetic regulators^56–58^. Our findings suggest that these systems could be more effective and less toxic if tested at lower light doses. Therefore, the relationship between a suitable light dosage and EL222 copy number contradicts the common perception that maximum light is always associated with maximum induction. In this study, we characterize the light dosage by intensity, but utilizing light pulses is also a suitable strategy for input delivery, as the tunability of our low copy yEGFP strain (Supplementary Fig. S7) and other studies suggest^29,33,34^.

The protective effect of *PMP20* suggests that the inhibitory effect of light on the growth and protein production of engineered strains stems from the emission of ROS by EL222, which is exacerbated when cells expressing high levels of this transcription factor are exposed to excessive light (Supplementary Fig. S10). This is consistent with previous studies on the generation of ROS by LOV-based proteins^46,47^. Pmp20p is an endogenous peroxiredoxin involved in degrading hydrogen peroxide and organic hydroperoxides at the peroxisome membrane surface^48^. While it is known to be upregulated in methanol and virtually absent in glycerol^42,59^, its activity in ROS-generating conditions without methanol has not been examined. ROS emitted by LOV-based proteins are generally singlet oxygen and, to a lesser degree, hydrogen peroxide^46^. This raises the possibility that Pmp20p is also capable of neutralizing singlet oxygen, or that EL222 emits hydrogen peroxide as the cause of its phototoxicity. In future studies, it could be interesting to test whether optimizing expression levels of *PMP20* in engineered strains would increase their tolerance to higher EL222 and blue light levels to further enhance protein production.

Understanding the link between EL222 concentration and phototoxicity can simultaneously enable optimization of light intensity throughout the process and facilitate the selection of the most suitable system for a given application. For strains requiring low copy numbers of the target protein, the simplicity of the coupled system makes it the preferred choice as it requires only one transformation. This strategy can be implemented by replacing yEGFP in the SMH131 plasmid (highlighted in Supplementary Table S1) with the gene of interest and integrating into *K. phaffii* using zeocin selection. However, in situations where high copy numbers of the gene of interest are preferred, the decoupled system is a better option to avoid the toxic effects of high copy numbers of EL222. To support adoption of this strategy, we produced a set of strains with a range of EL222 copy numbers that can be used as parent strains (highlighted in Supplementary Table S2) in which to integrate the gene of interest under the control of P_C120_ at varying and decoupled copy numbers (Supplementary Fig. S3). This can be achieved by replacing yEGFP in plasmid SMH234 or Mfp5 in SMH244 (highlighted in Supplementary Table S1) with the gene of interest, depending on whether the protein is meant to be secreted or produced intracellularly, and using the resulting plasmid to transform the parent strains with zeocin selection. For applications where light penetration may be a concern (for example in large-scale bioreactors), ySMH203-2 and ySMH203-21 (containing 5 and 8 copies of EL222, respectively) may be the best strains to use, as they are more responsive to lower light levels (highlighted in Supplementary Table S2). In cases where light penetration is not a concern, strain ySMH203-16 (housing only 1 copy of EL222) offers a more forgiving range of suitable light intensities, making it easier to avoid phototoxicity (highlighted in Supplementary Table S2).

The hybrid dynamic model for yEGFP production, influenced by the EL222 copy number and light intensity (Fig. 3), outlines a suitable strategy for capturing the dynamic behavior of optogenetic bioprocesses for recombinant protein production. This modeling strategy holds the potential to unlock advanced open-loop and closed-loop model-based optimization schemes in future studies. Optimal control problems could be formulated to maximize bioprocess performance^60–63^, exploiting the available degrees of freedom while subject to the constraints imposed by some knowledge of the system (in this case, the hybrid dynamic model), as well as potential economic, technical, safety, and environmental considerations. In the context of optogenetic recombinant protein production, degrees of freedom of interest include variables such as inoculum size, initial substrate concentration, strain selection with a specific EL222 copy number, dynamic light intensity trajectories as in two-phase processes, and feed rates as in fed-batch bioreactors. Furthermore, the modeling strategy enables the potential use of Bayesian-related optimizations^64^. Incorporating the variance of the Gaussian processes into model-based optimization problems could reveal areas with high (parameter) uncertainty, facilitating the balancing of the system’s exploration and exploitation^65^. The implementation of these advanced model-based optimization and control strategies in optogenetically assisted bioprocessing is currently under development in our group.

When applying the decoupled system to produce secreted proteins, an unexpected finding was that production in glucose is weaker than in glycerol and, in most cases, methanol. This was observed for all three protein targets tested (yEGFP, β-lactoglobulin, SR18) and at both low and high EL222 copy numbers (Fig. 6). Though the mechanism causing this difference is not understood, we hypothesize that it is related to some interference between glucose, light, and the secretion pathway, as intracellular protein production levels using light induction are similar between glucose and glycerol (Supplementary Fig. S5). Under our tested conditions, we found that glycerol is the best carbon source for optogenetic control of protein secretion, including to obtain superiority to methanol induction, which informed our substrate of choice when scaling to the bioreactor level. Future studies will be necessary to elucidate the mechanism by which protein secretion is diminished when using light induction in glucose media to overcome this limitation.

Our bioreactor experiments show that light induction of protein secretion can be successfully achieved in 1-L cultures at very high cell densities, even with a strain containing only one copy of EL222 (Fig. 7, Supplementary Fig. S22 and S23). Using this strain provided us with a wider range of suitable light conditions, which facilitated the scaleup process. However, this strain also shows the lowest light responsiveness of the parent strains we developed, leaving open the possibility of examining different light doses at this scale. These results also raise the prospect of obtaining robust light induction in larger pilot- or demonstration-scale bioreactors using strains with higher copy numbers of EL222, which are at least 10 times more sensitive to light (based on observations that strains with one copy need 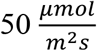 to maximize induction, while strains containing eight copies are fully active at only 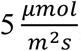, and possibly less) (Fig. 3). Further scaleup of light-induced protein production in *K. phaffii* can be conducted in the future by adapting existing industrial photobioreactors used in microalgae and cyanobacteria processes, or by applying known photobioreactor design principles to create new systems^66–70^. Altogether, our results reduce concerns of light penetration in lab-scale bioreactors and bode well for future development of optogenetically controlled industrial-scale processes.

## Conclusions

The optogenetic systems and methods we developed demonstrate that light is a suitable inducer or control input in the context of dynamic metabolic control and recombinant protein production by *K. phaffii*. As such, our work offers a promising alternative to conventional methanol induction, with the potential to enhance process tunability and flexibility. The interplay between EL222 copy number and light dosage emerges as a critical degree of freedom for future optimization and control, particularly in model-based approaches and bioreactor design. Our proposed methods augment the toolbox of metabolic and bioprocess engineering for designing and operating recombinant protein production processes. Moreover, beyond the outlined applications, this work lays the groundwork for advanced cybergenetic approaches to dynamic metabolic control using optogenetics, including feedback control strategies such as model predictive control and reinforcement learning.

## Supporting information

Supplemental Figures, Tables, and Sequences

## Acknowledgements

This research was supported by the U.S Department of Energy, Office of Science, Office of Biological and Environmental Research Award Number DE-SC0022155, and NSF Award MCB-2300239 (to JLA), as well as the Maeder Graduate Fellowship in Energy and the Environment (to SMH), and NSF grant number DGE-2039656 (to SKK).

## Author contributions

SMH and JLA conceived the project; SMH, MAL, SKK, and JLA designed and analyzed the experiments; SMH, MAL, and SKK executed the experiments; SER performed the modeling, SMH, SER, SKK, and JLA wrote the paper; JLA secured funding and supervised the project. All authors read and approved the final manuscript.

## Materials and data availability

Correspondence and requests for materials and data should be addressed to José L. Avalos

## Code availability

The code related to the Gaussian-process-supported hybrid model is available in the following repository: https://bitbucket.org/codes-to-share/gp_model_sh/src/main/

