## Supplemental Figures, Tables, and Sequences for "Balancing doses of EL222 and light improves optogenetic induction of protein production in *Komagataella phaffii*"

[1] Department of Chemical and Biological Engineering, Hoyt Laboratory, Princeton University, Princeton, NJ, USA; [2]The Omenn-Darling Bioengineering Institute, Princeton University, Princeton, NJ, USA; [3]The Andlinger Center for Energy and the Environment, Princeton University, Princeton, NJ, USA; [4]Department of Molecular Biology, Princeton University, Princeton NJ 08544, USA; [5]High Meadows Environmental Institute, Princeton University, Princeton NJ 08544, USA. [+]Current affiliation: Commonwealth Scientific and Industrial Research Organisation, Clayton, VIC, 3168, AUS.

[‡]Co-first authorship.

*Corresponding author: José L. Avalos

Department of Chemical and Biological Engineering,

Princeton University,

101 Hoyt Laboratory, William Street, Princeton, NJ 08544, USA

Phone lab: +1 (609) 258-0542

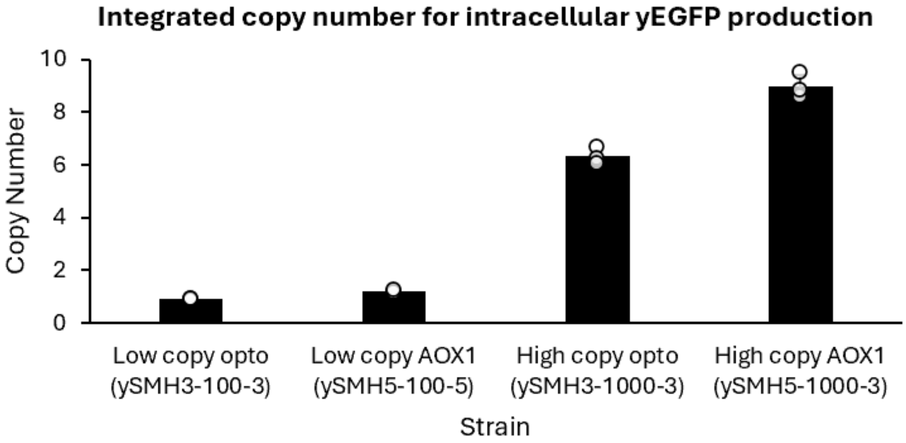


**Supplementary Figure S1. Copy number analysis of the intracellular yEGFP-producing strains.** Optogenetic and methanol-induced strains were analyzed by qPCR to identify single copy strains and comparable high-copy transformants. Single copy strains were isolated from plates containing 100 µg/mL Zeocin, while the high copy strains were identified on 1000 µg/mL Zeocin plates. Data is shown as mean values and errors bars depict the standard deviation of three independent replicates.

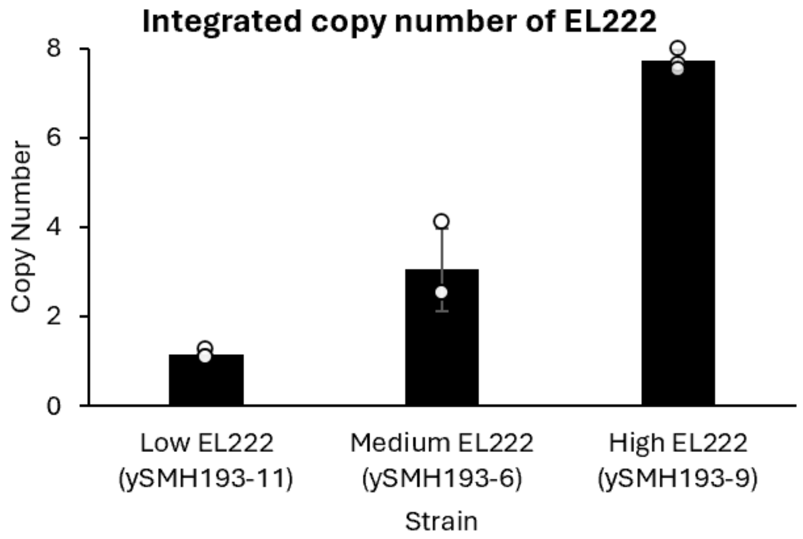


**Supplementary Figure S2. qPCR analysis for strains used to characterize the effect of EL222 copy number on light sensitivity.** Copy numbers are shown for the three strains used to examine the impact of EL222 amount on strain sensitivity. Data is shown as mean values and errors bars depict the standard deviation of three independent replicates.

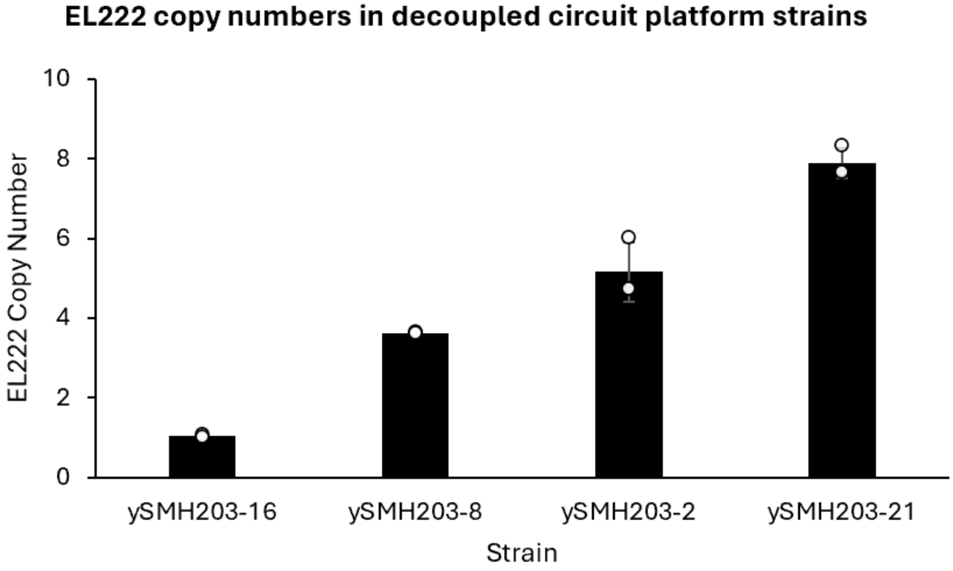


**Supplementary Figure S3. qPCR analysis of EL222-containing strains for the decoupled optogenetic system.** qPCR was used to identify strains containing a diverse range of EL222 copy numbers ranging from 1 to 8. These strains served as parents for subsequent experiments which used the decoupled optogenetic system. Data is shown as mean values and error bars depict the standard deviation of three independent replicates.

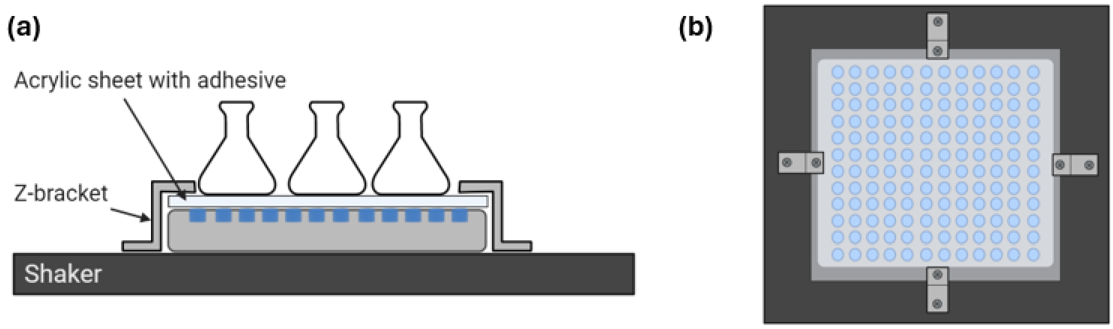


**Supplementary Figure S4. Shake flask illuminator platform.** An apparatus constructed from widely available materials enabled homogenous illumination of shake flasks, which is shown from an (a) side and (b) top view. Details regarding the construction of this apparatus can be found in the "Illumination and measurement of light intensity" section of the Methods.

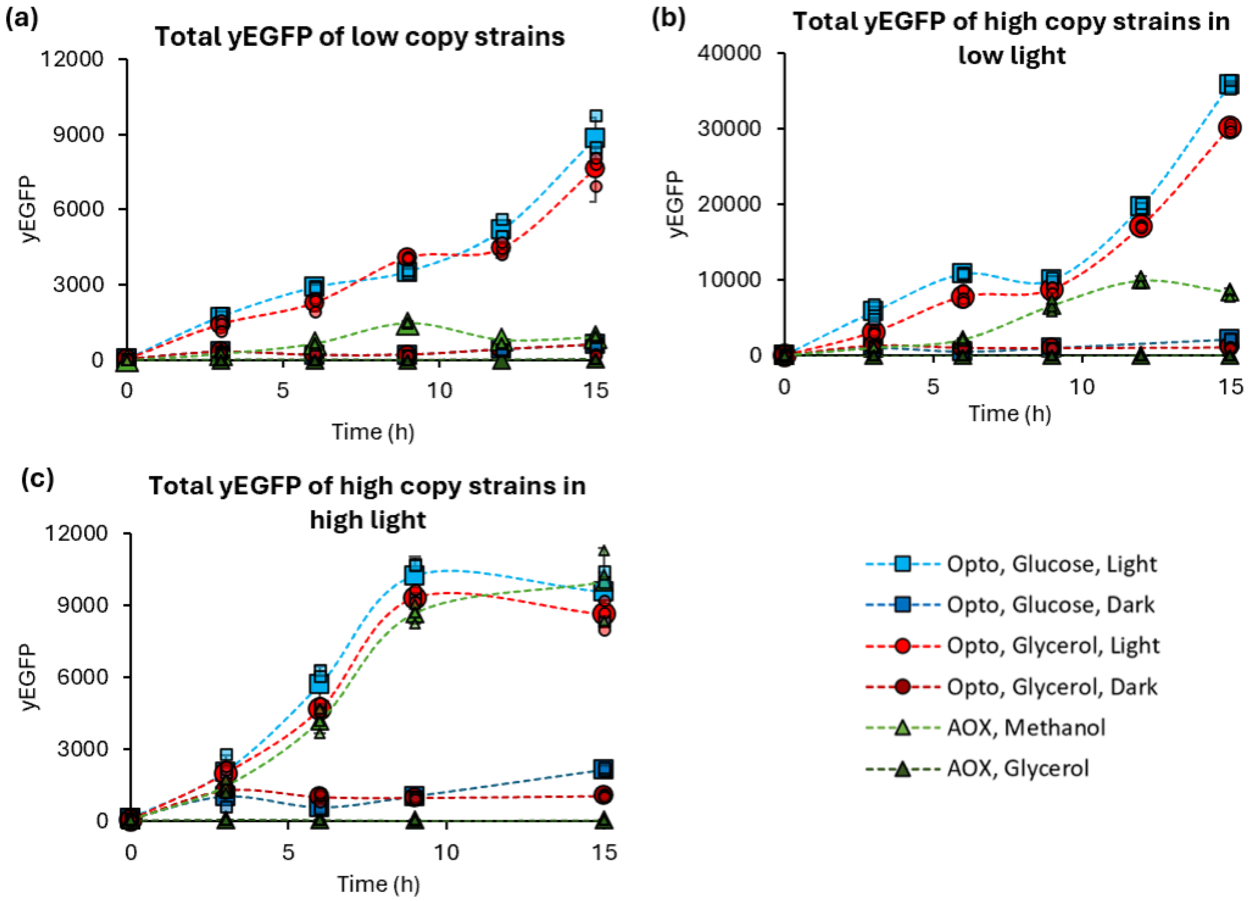


**Supplementary Figure S5. Total fluorescence with the coupled optogenetic system.** Total production of yEGFP is compared between the coupled optogenetic system and $P_{AOX1}$ at (a) a single copy in 70 $\frac{\mu mol}{m^2 s}$ of light as well as at ~8 copies in (b) 5 $\frac{\mu mol}{m^2 s}$ and (c) 70 $\frac{\mu mol}{m^2 s}$ light intensities. Data is shown as mean values, with the smaller icons representing each individual replicate. Error bars depict the standard deviation of three independent biological replicates.

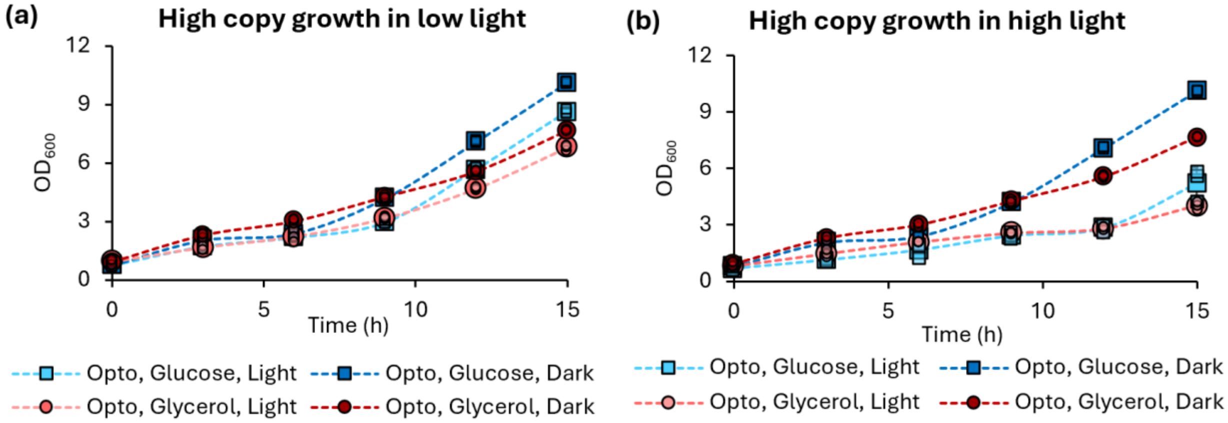


**Supplementary Figure S6. Impact of light on growth in an optogenetic yEGFP-producing strain.** Growth in the strain containing a high copy of the 'coupled' optogenetic system (ySMH3-1000-3) is shown at (a) a low light intensity of 5 $\frac{\mu mol}{m^2 s}$ and (b) a higher light intensity of 70 $\frac{\mu mol}{m^2 s}$ in BMD1 and BMG1 media. The effect on growth is minimal in the low intensity condition, but severe at 70 $\frac{\mu mol}{m^2 s}$. Data is shown as mean values with smaller icons representing each of the three independent biological replicates.

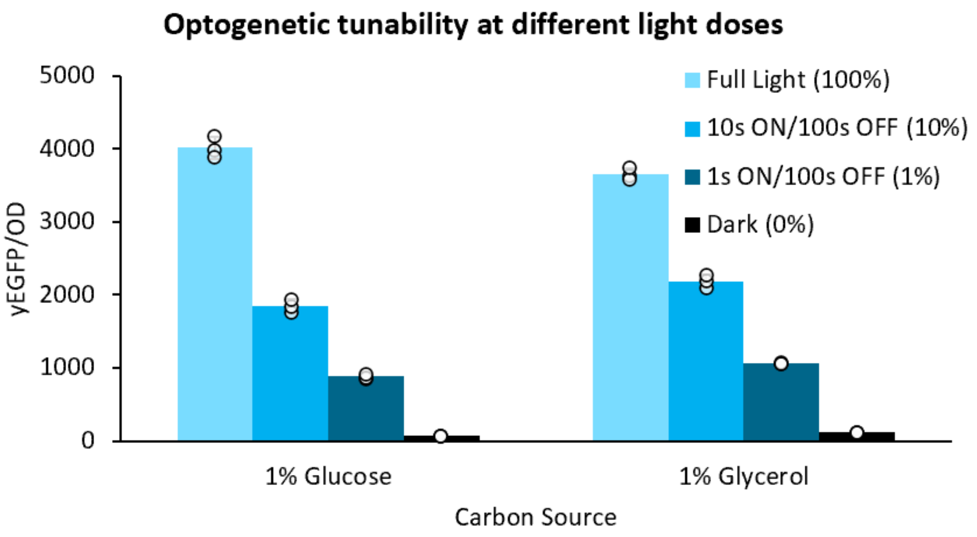


**Supplementary Figure S7. Tunability of optogenetic expression by light pulsing.** Optogenetic control offers new capabilities relative to methanol induction because expression levels can be tuned by adjusting the light schedule. Light dosage was mediated through pulsing light of 70 $\frac{\mu mol}{m^2 s}$ intensity at different schedules over ySMH3-100-3 cells in BMD1 and BMG1 media. Data is shown as mean values and errors bars depict the standard deviation of three independent replicates.

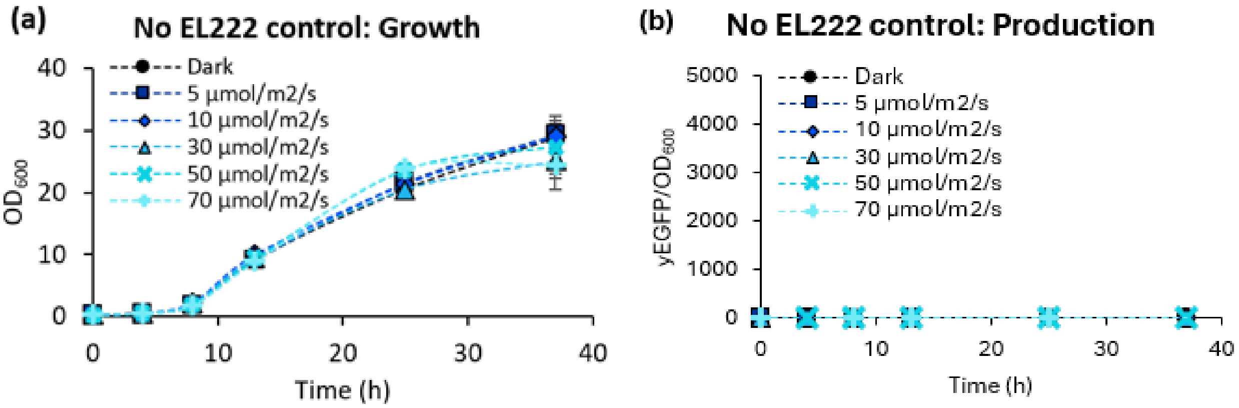


**Supplementary Figure S8. Effect of light dosage on a control strain without EL222.** (a) Growth and (b) production of yEGFP were analyzed in a strain containing $P_{C120}$-yEGFP but no EL222 (ySMH115-14). Without EL222, light has no effect on growth and there is no production from the $P_{C120}$ promoter. Data is shown as mean values and error bars depict the standard deviation of three independent replicates.

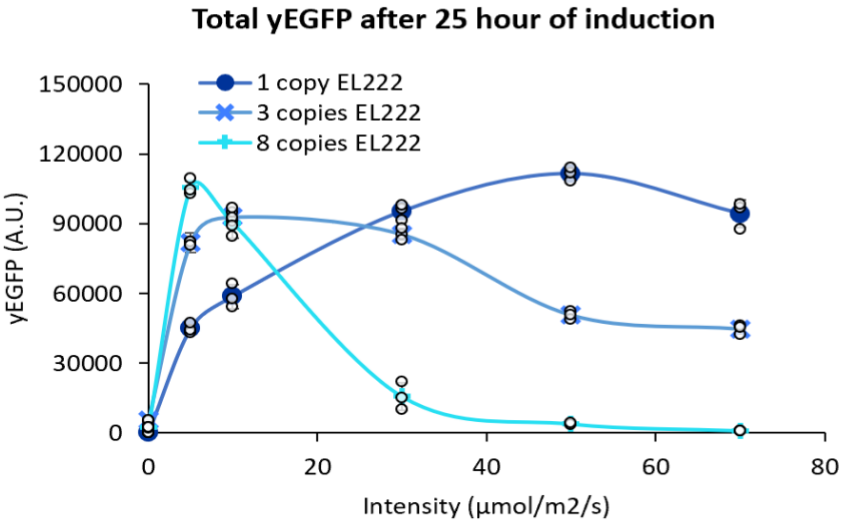


**Supplementary Figure S9. Optimizing light dosage based on EL222 copy number.** The total yEGFP (not normalized by cell density) after 25 hours is shown for strains containing 1, 3, and 8 copies of EL222 to depict the optimal light dosage for each strain. Data is derived from the same experiment as Figure 3 with points showing the average of three independent biological replicates, and error bars representing the standard deviation.

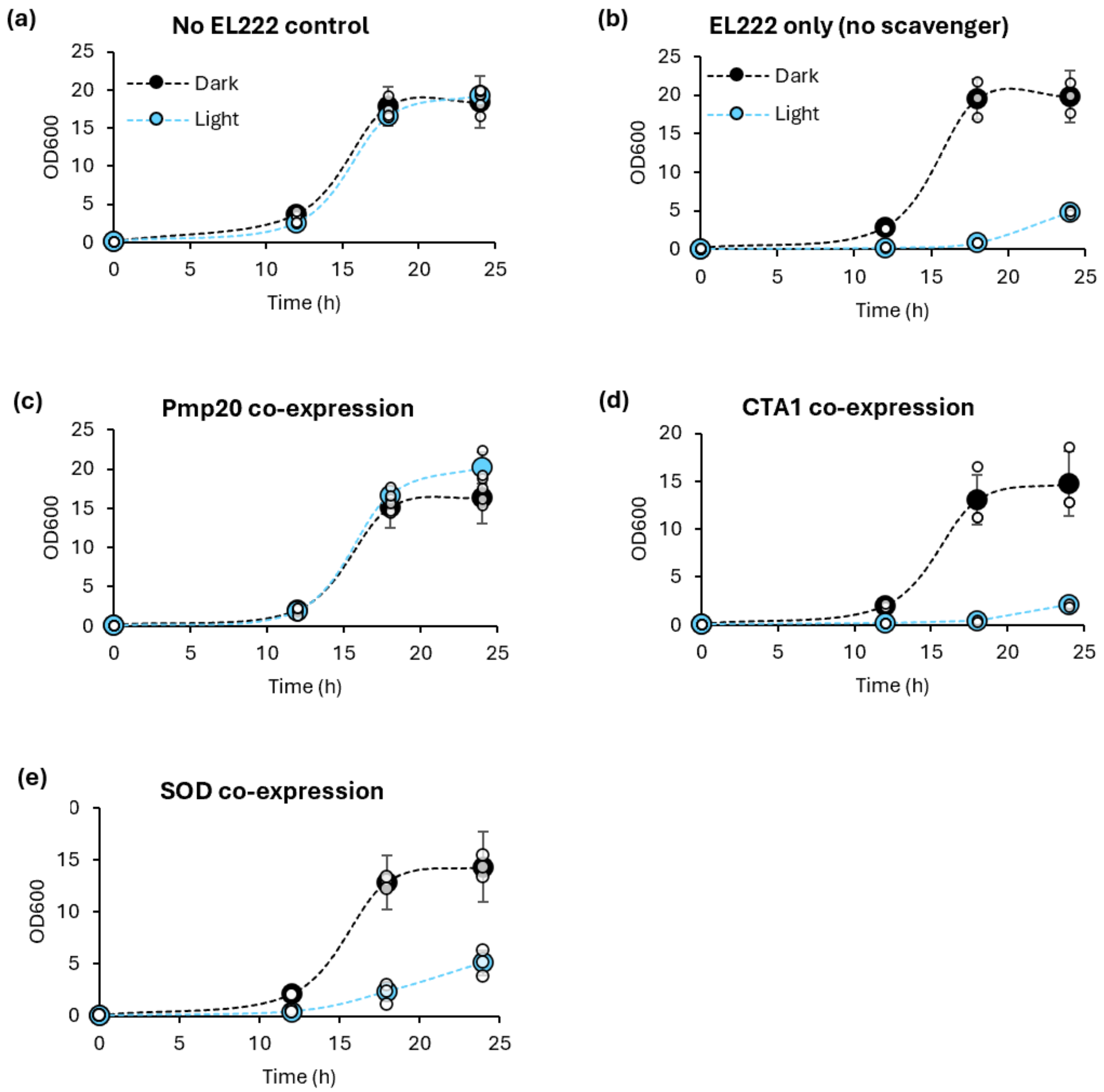


**Supplementary Figure S10. Co-expression of *K. phaffii* ROS scavengers in strains experiencing phototoxicity.** Three ROS scavengers native to *K. phaffii* were co-expressed with EL222 and tested for recovery of growth in the light at 70 $\frac{\mu mol}{m^2 s}$ in 1% glucose medium. (a) A control strain without EL222 (ySMH196) is unaffected by light, while (b) a strain containing EL222 (ySMH197) exhibits severe toxic effects. Co-expression of (c) *PMP20* (ySMH210) neutralizes the phototoxic effect while expression of the (d*) CTA1* catalase (ySMH207) or (e) *SOD1* superoxide dismutase (ySMH209) does not recover growth. Data is shown as mean values and errors bars depict the standard deviation of three independent replicates.

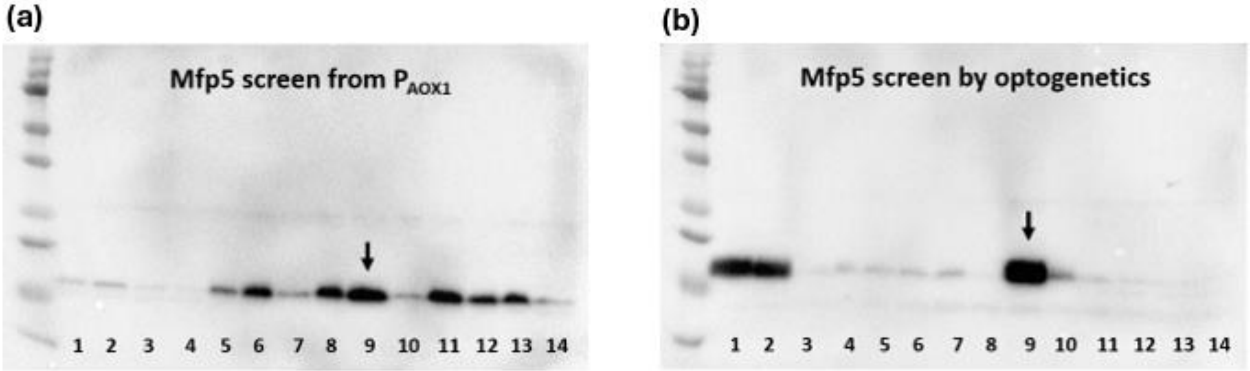


**Supplementary Figure S11. Screening for the best Mfp5-producing strains.** Fourteen (a) $P_{AOX1}$-driven and (b) optogenetic Mfp5-producing strains were screened and analyzed with western blot to identify the best producers to later compare in triplicate. $P_{AOX1}$-controlled strains (ySMH186) were induced for 48 hours in BMM1 medium, while the optogenetic strains (ySMH235) were induced for 48 hours in BMD1 medium and light at 50 $\frac{\mu mol}{m^2 s}$ intensity. Each lane represents a different colony tested, and arrows indicate the best-performing strain that was selected for subsequent experiments.

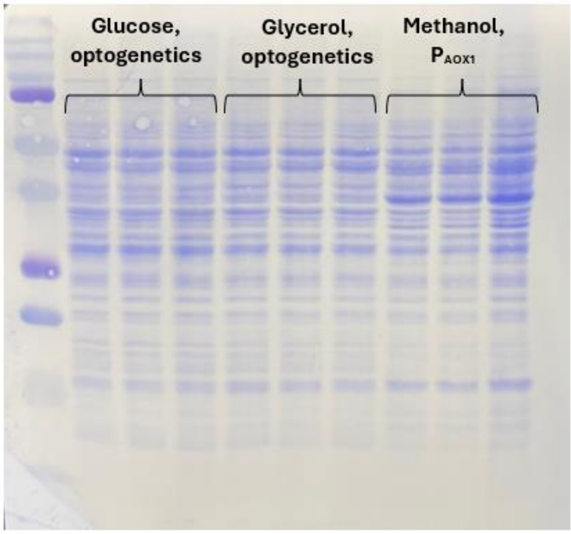


**Supplementary Figure S12. Equal loading verification for the Mfp5-production test.** After completing the western blot comparing optogenetic and methanol-induced intracellular production of Mfp5, a Coomassie stain was performed on the membrane to verify comparable loading between wells.

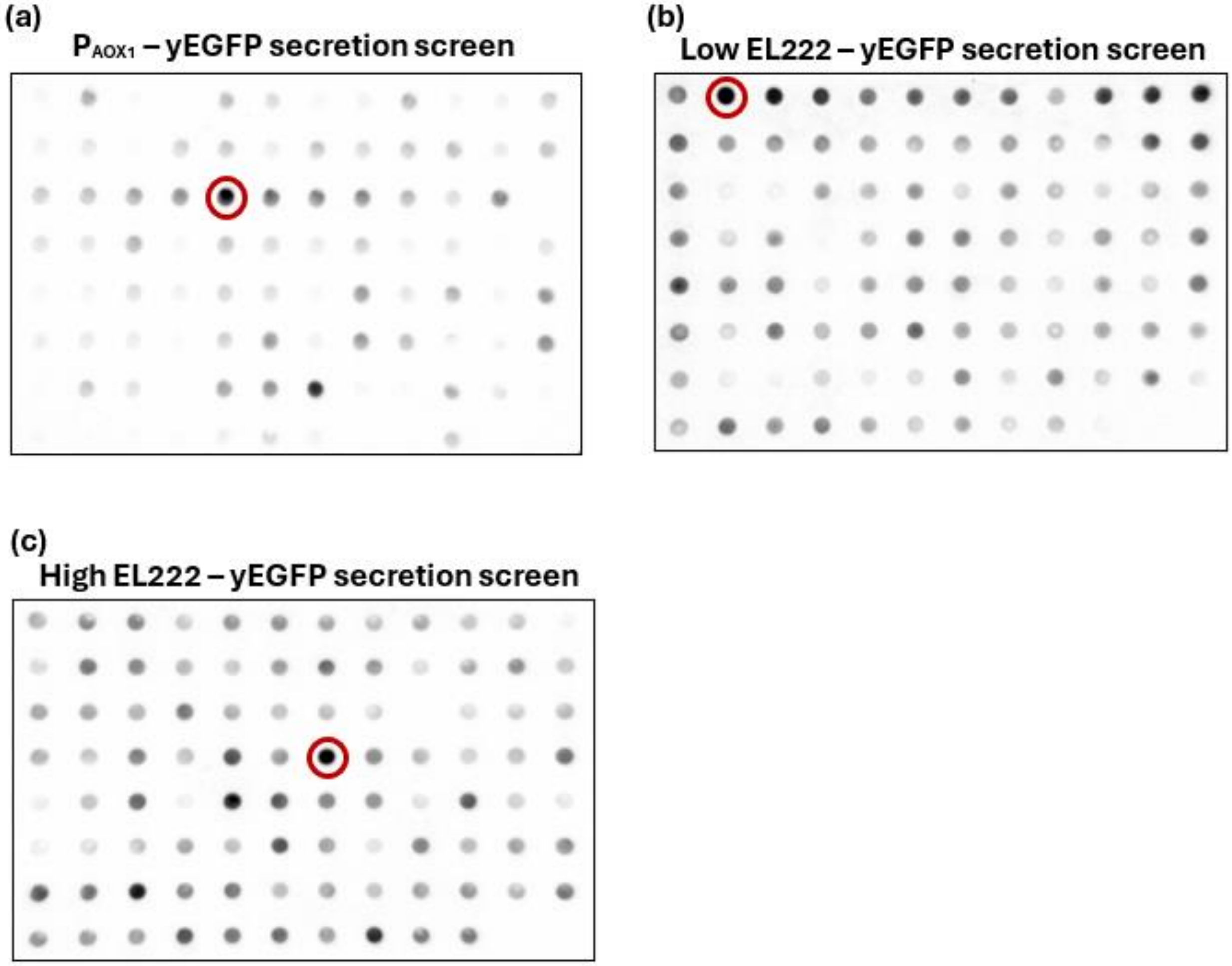


**Supplementary Figure S13. Screening for the best yEGFP-secreting strains.** yEGFP-secreting strains induced from either (a) $P_{AOX1}$ (ySMH154) or the decoupled optogenetic system with (b) one (ySMH217) or (c) five copies of EL222 (ySMH219) were screened in a 96-well format using a dot blot analysis. All strains were induced for 48 hours with in either (a) BMM1 or (b, c) BMD1 medium. Circled dots indicate the best producers, which were then used in downstream experiments.

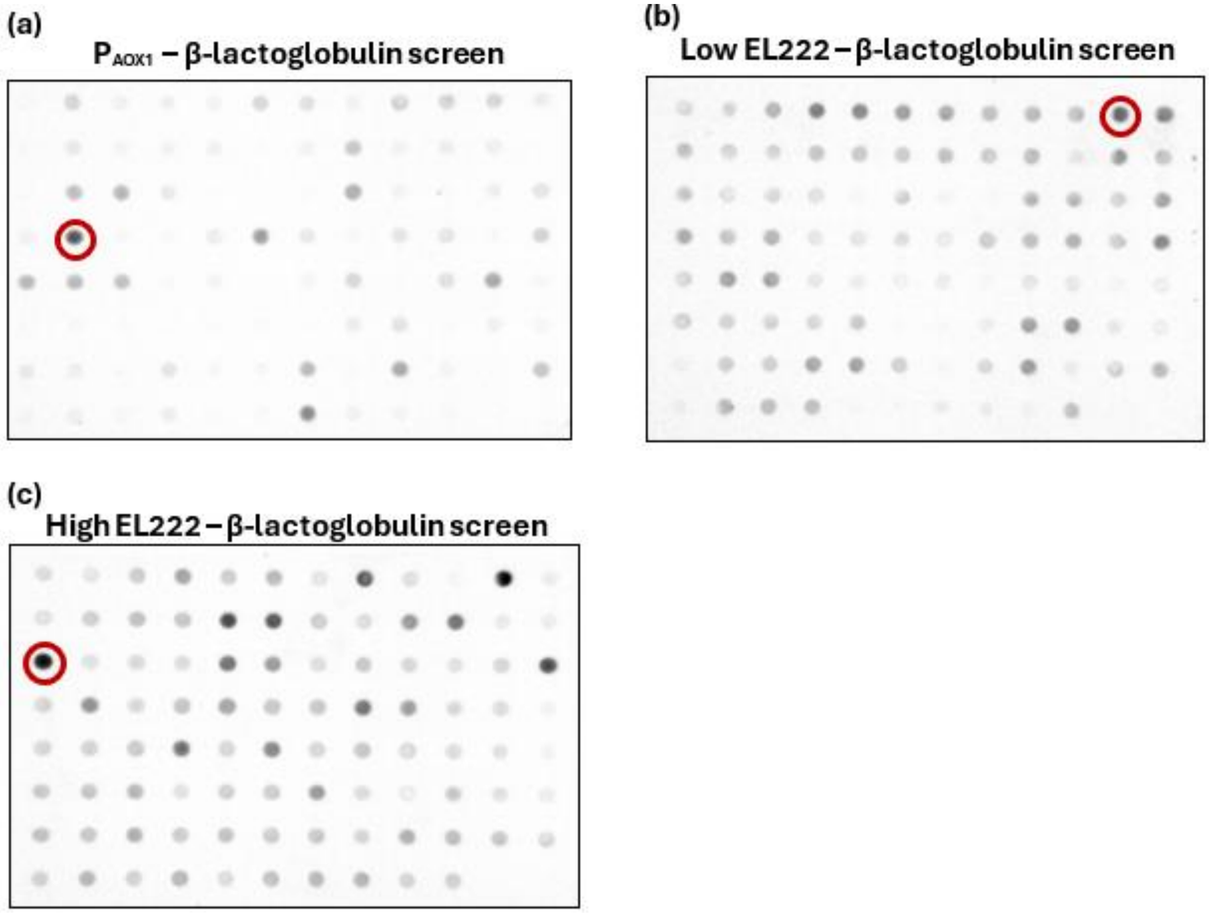


**Supplementary Figure S14. Screening for the best β-lactoglobulin secreting strains.** β-lactoglobulin production was screened in a 96-well dot blot format with either (a) $P_{AOX1}$ (ySMH51) or the decoupled optogenetic system with (b) one or (c) five copies of EL222 (ySMH224 and ySMH226). All strains were induced for 48 hours in either (a) BMM1 or (b, c) BMD1 medium. Circled dots indicate the best producers, which were then used in downstream experiments.

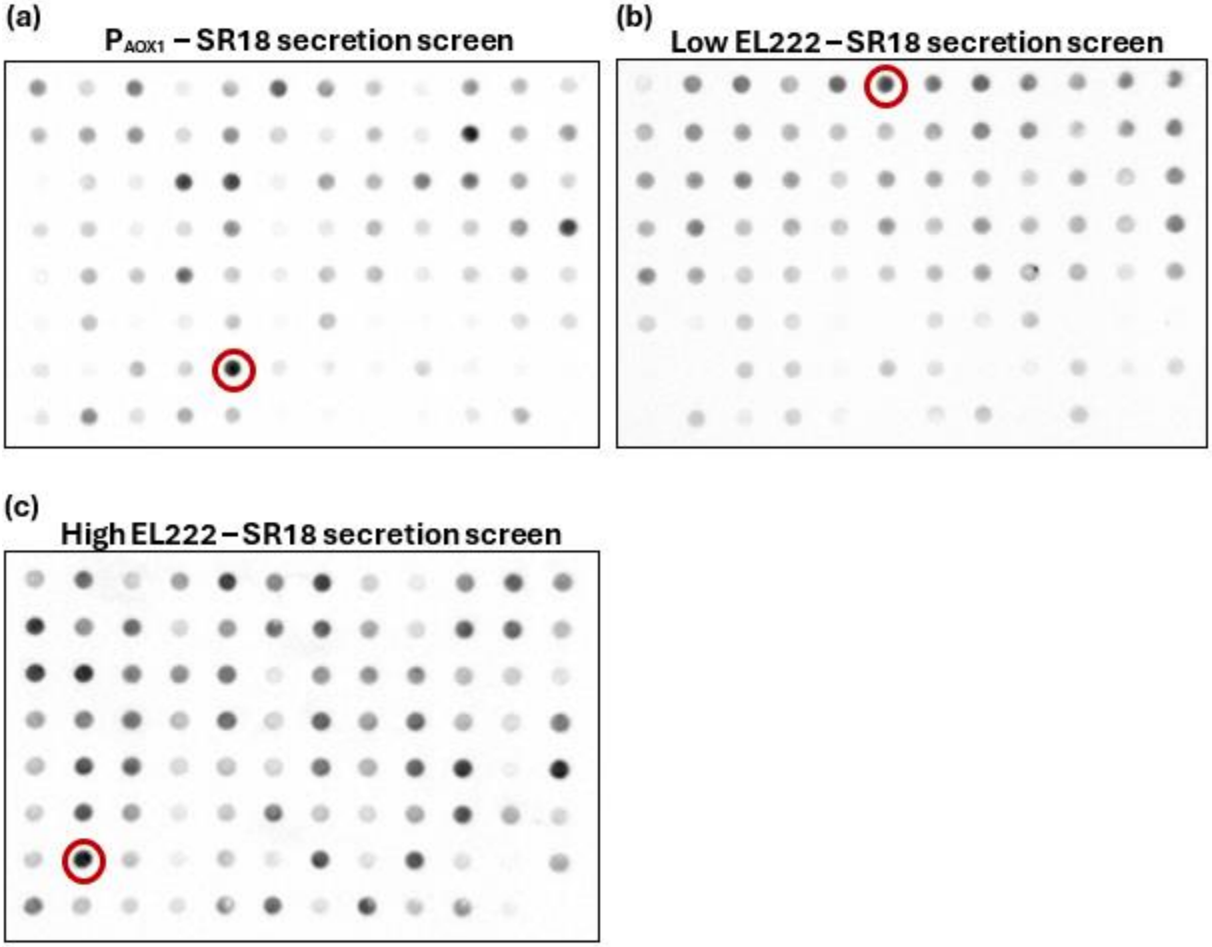


**Supplementary Figure S15. Screening for the best SR18 nanobody secreting strains.** Production of the SR18 nanobody was screened in a 96-well dot blot format from either (a) $P_{AOX1}$ (ySMH223) or the decoupled optogenetic system with (b) one or (c) five copies of EL222 (ySMH230 and ySMH231). All strains were induced for 48 hours in either (a) BMM1 or (b, c) BMD1 medium. Circled dots indicate the best producers, which were then used in downstream experiments.

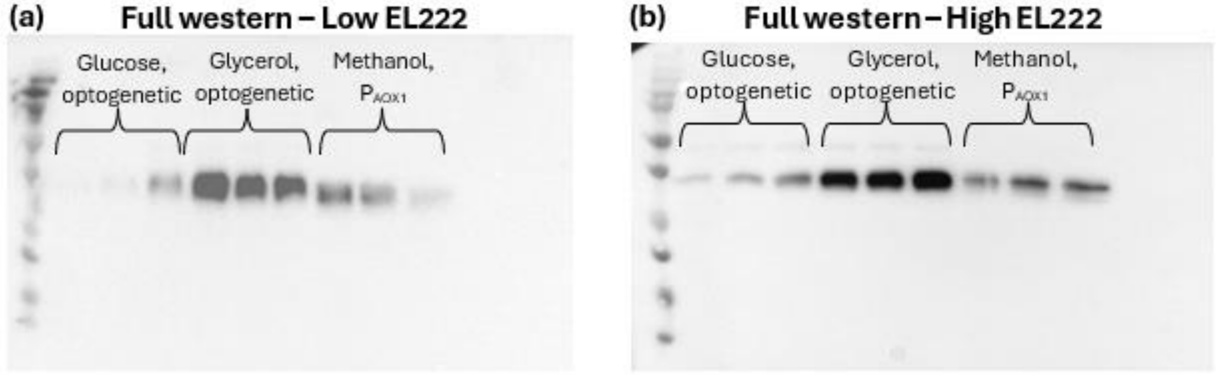


**Supplementary Figure S16. Uncropped western blots of secreted yEGFP production.** Full images of the western blots comparing the (a) low EL22 and (b) high copy EL222 yEGFP-secreting strains (ySMH217-A2 and ySMH219-D7) against the best $P_{AOX1}$-driven producer (ySMH154-C5), which serve as the basis to Figure 6 a-b.

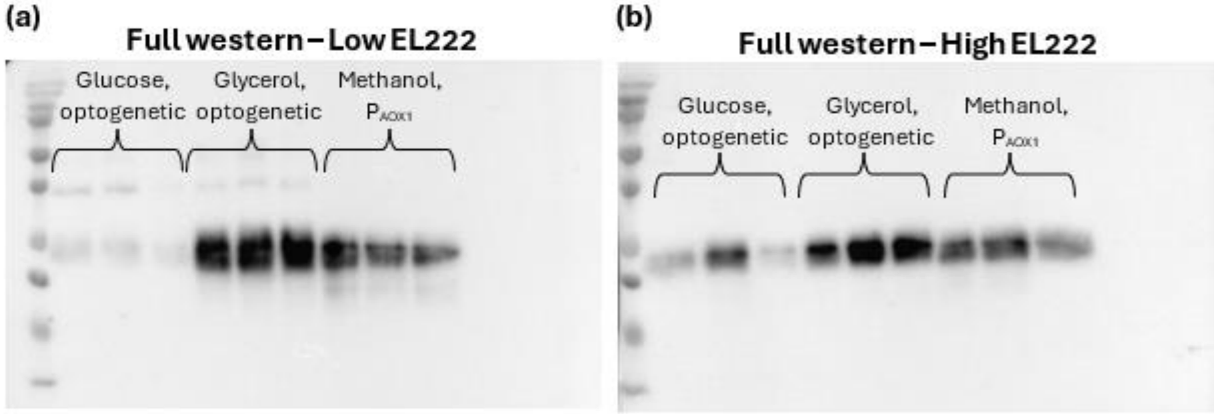


**Supplementary Figure S17. Uncropped western blots of secreted β-lactoglobulin production.** Full images of the western blots comparing the (a) low EL222 and (b) high copy EL222 β-lactoglobulin secreting strains (ySMH224-A11 and ySMH226-C1) against the best $P_{AOX1}$-driven producer (ySMH51-D2), which serve as the basis to Figure 6 c-d.

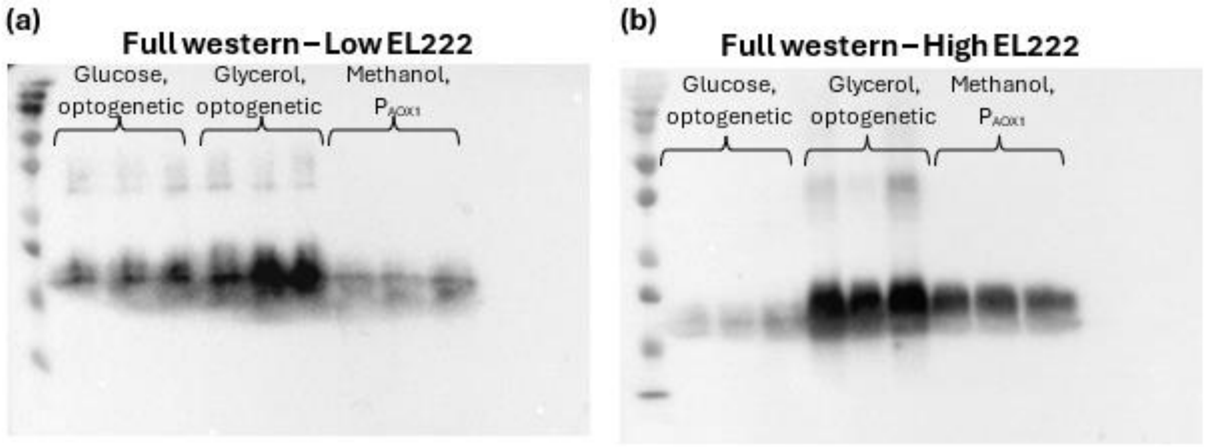


**Supplementary Figure S18. Uncropped western blots of secreted SR18 nanobody production.** Full images of the western blots comparing the (a) low EL222 and (b) high copy EL222 SR18-secreting strains (ySMH230-A6 and ySMH231-G2) against the best $P_{AOX1}$-driven producer (ySMH223-G5), which serve as the basis to Figure 6 e-f.

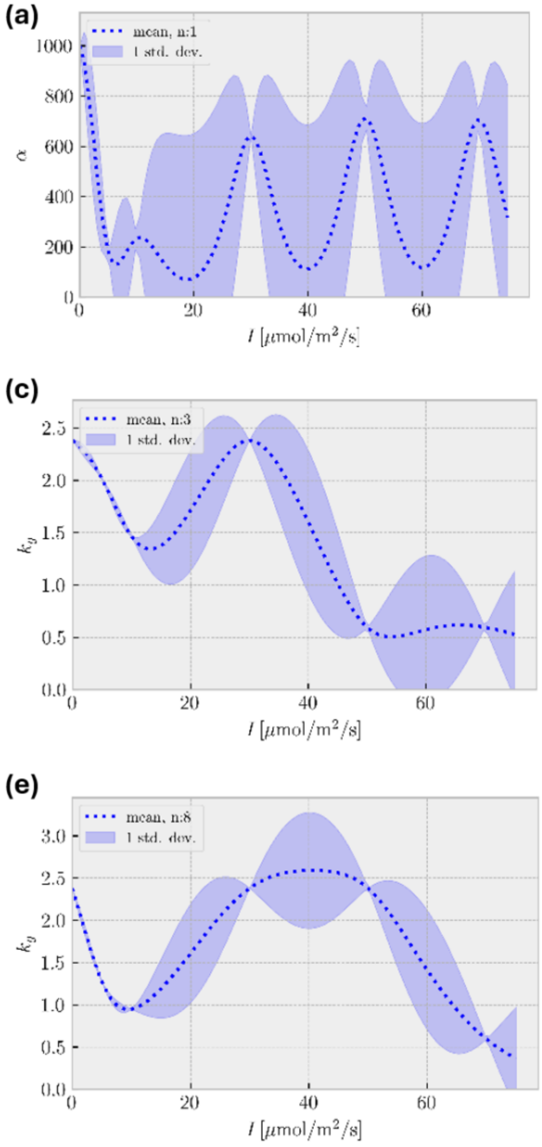


**Supplementary Figure S19.** Selected Gaussian-process-predicted parameters at different light intensities and copy numbers for the hybrid model in Figure 3. We depict functions for parameters $k_g$ (a, c, e) and $\alpha$ (b, d, f) based on light intensity, *I*, and EL222 copy number. Parameter functions for (a, b) one, (c, d) three, and (e, f) eight copies of EL222 are shown from light intensities ranging from total darkness to 70 $\frac{\mu mol}{m^2 s}$. The predicted mean is indicated by dotted blue lines and the predicted standard deviation by a blue shadowed area.

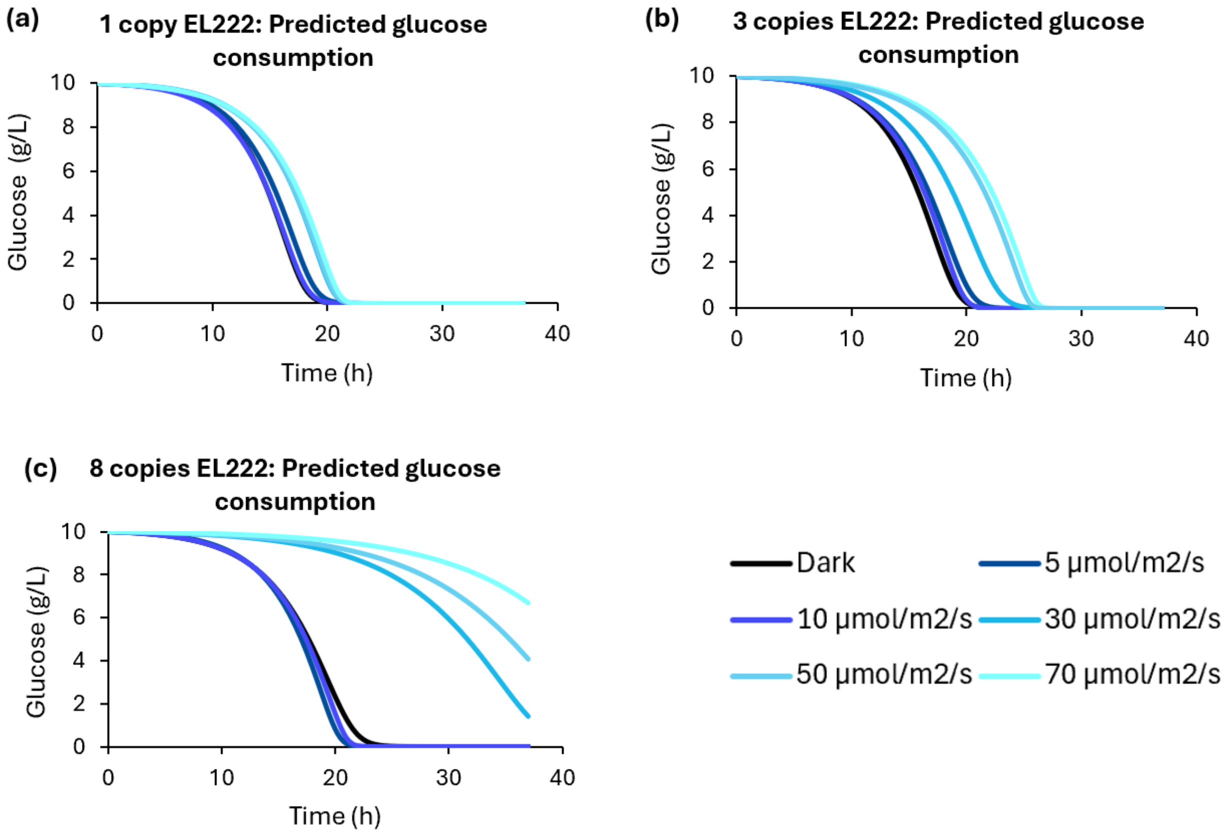


**Supplementary Figure S20. Model-predicted glucose concentrations for the experiments shown in Figure 3.** Predictions according to the hybrid Gaussian-process-supported model are shown throughout time for each light intensity tested with (a) one, (b) three, and (c) eight copies of EL222.

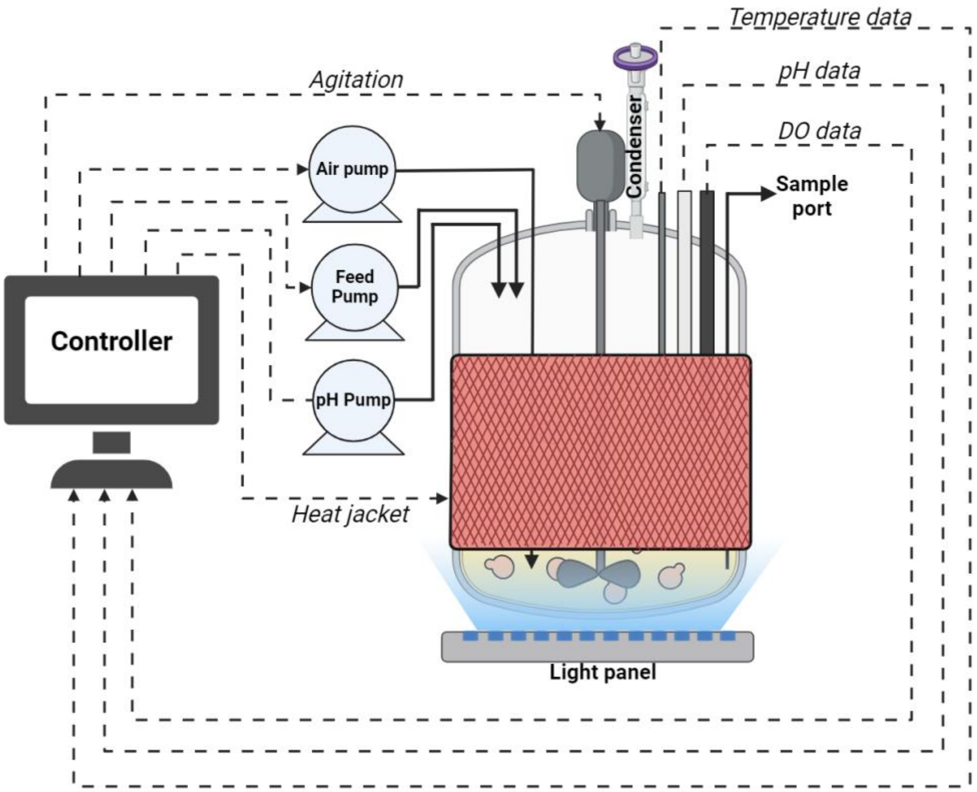


**Supplementary Figure S21. Light-controlled bioreactor schematic.** The optogenetic culture was illuminated from the bottom of the culture using a light panel placed below the reactor. Temperature, pH, and dissolved oxygen (DO) probes communicated to a controller which adjusted the agitation and air flow (DO control), addition of ammonium hydroxide (pH control), or heating (temperature control) as needed.

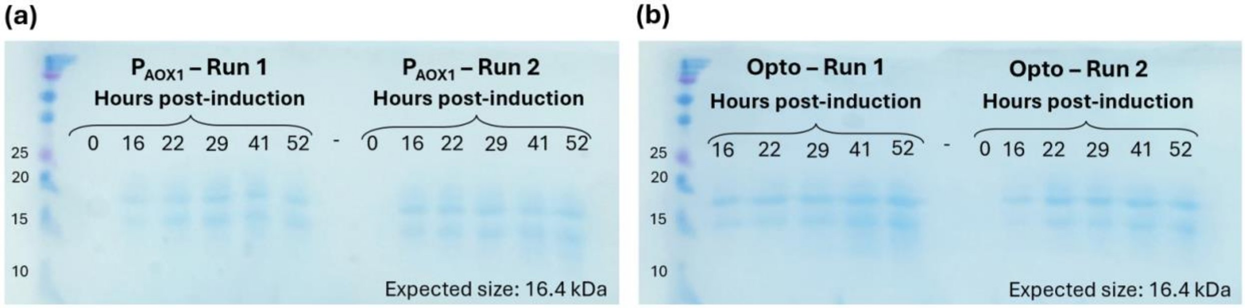


**Supplementary Figure S22. SR18 production in lab-scale bioreactor runs.** Samples from two replicate bioreactor runs of (a) methanol-induced and (b) light-induced SR18 production analyzed by SDS-PAGE with Coomassie staining. The samples were obtained at the indicated time points after induction, and gels were run, stained, and destained at the same time. Lanes marked with a negative sign (-) represent blank wells as a negative control. A 0-hour timepoint sample was not obtained for the first light-induced bioreactor run, so analysis for this timepoint is based on the second run performed.

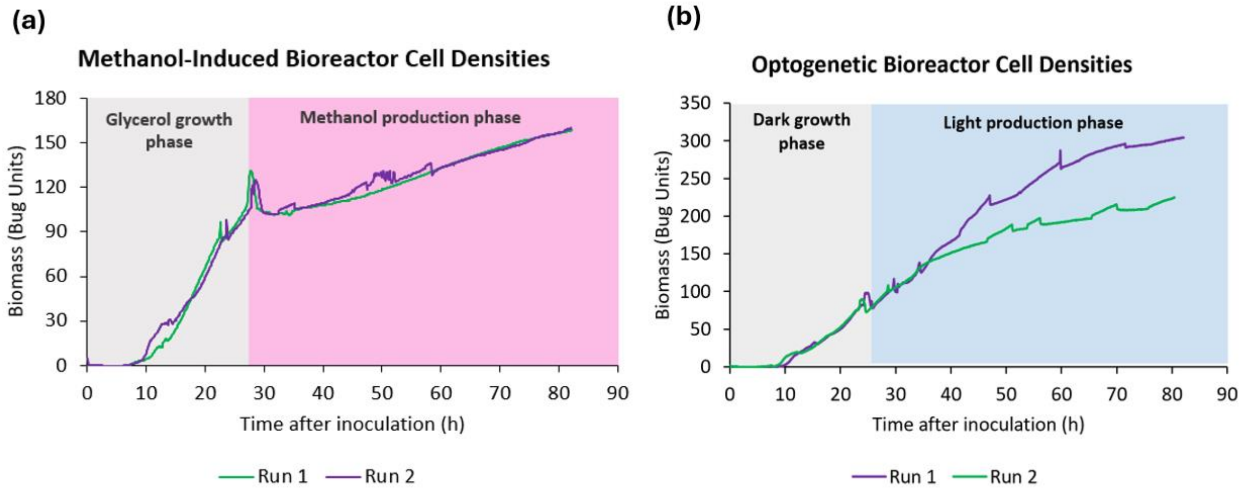


**Supplementary Figure S23. Cell densities in the bioreactor experiments.** Biomass measurements, quantified as the raw Bug Unit values, during the (a) methanol-induced and (b) optogenetic bioprocesses are shown, with the growth and production phases of the processes annotated.

**Supplementary Table 1. Plasmids used in this study.**

| Plasmid | Purpose | Description | Marker | Integration locus (Cut site) | Source |
|---|---|---|---|---|---|
| pPICZαA | Parent plasmid | $P_{AOX1}$-$\alpha_{sec}$-$TT_{AOX1}$ | Zeocin | $P_{AOX1}$ (PmeI) | Invitrogen |
| pCri15b | Parent plasmid | $P_{AOX1}$-EYFP-$TT_{AOX1}$ | Zeocin | $P_{AOX1}$ (PmeI) | Goulas et al.[1] |
| A171 | CRISPR parent plasmid | $P_{HTX1}$-Cas9-$TT_{DAS1}$, $P_{HTX1}$-gRNA-$TT_{AOX1}$ | Zeocin | N/A | Bisy GmbH |
| SMH5 | Intracellular yEGFP from $P_{AOX1}$ | $P_{AOX1}$-yEGFP-$TT_{AOX1}$ | Zeocin | $P_{AOX1}$ (PmeI) | This study |
| SMH6* | Intracellular yEGFP from $P_{C120}$ (no EL222) | $P_{C120}$-yEGFP-$TT_{AOX1}$ | Zeocin | $P_{AOX1}$ (PmeI) | This study |
| SMH60 | *HIS4* CRISPR knockout | $P_{HTX1}$-Cas9-$TT_{DAS1}$, $P_{HTX1}$-gRNA$_{HIS4}$-$TT_{AOX1}$ | Zeocin | N/A | This study |
| SMH71 | β-lactoglobulin secretion from $P_{AOX1}$ | $P_{AOX1}$-$\alpha_{sec}$-$\beta_{lacto}$-$TT_{AOX1}$ | Zeocin | $P_{AOX1}$ (PmeI) | This study |
| SMH131* | Intracellular yEGFP from coupled system | $P_{C120}$-yEGFP-$TT_{AOX1}$, $P_{ADH2}$-EL222-$TT_{ADH2}$ | Zeocin | $P_{AOX1}$ (PmeI) | This study |
| SMH139 | EL222 for toxicity tests/decoupled system | $P_{ADH2}$-EL222-$TT_{ADH2}$ | Kan (*E. coli*) G418 (yeast) | $P_{ADH2}$ (ApaI) | This study |
| SMH158 | yEGFP secretion from $P_{AOX1}$ | $P_{AOX1}$-$\alpha_{sec}$-yEGFP-$TT_{AOX1}$ | Zeocin | $P_{AOX1}$ (PmeI) | This study |
| SMH207 | Intracellular Mfp5 from $P_{AOX1}$ | $P_{AOX1}$-Mfp5-$TT_{AOX1}$ | Zeocin | $P_{AOX1}$ (PmeI) | This study |
| SMH219 | *CTA1* scavenger co-expression | $P_{GAP}$-*CTA1*- $TT_{AOX1}$ | Amp (*E. coli*) *HIS4* (yeast) | *HIS4* (SalI) | This study |
| SMH229 | *SOD1* scavenger co-expression | $P_{GAP}$-*SOD1*- $TT_{AOX1}$ | Amp (*E. coli*) *HIS4* (yeast) | *HIS4* (SalI) | This study |
| SMH230 | *PMP20* scavenger co-expression | $P_{GAP}$-*PMP20*- $TT_{AOX1}$ | Amp (*E. coli*) *HIS4* (yeast) | *HIS4* (SalI) | This study |
| SMH234* | yEGFP secretion from decoupled system | $P_{C120}$- $\alpha_{sec}$-yEGFP-$TT_{AOX1}$ | Zeocin | $P_{AOX1}$ (PmeI) | This study |
| SMH240* | β-lactoglobulin secretion from decoupled system | $P_{C120}$- $\alpha_{sec}$- $\beta_{lacto}$-$TT_{AOX1}$, | Zeocin | $P_{AOX1}$ (PmeI) | This study |
| SMH242 | SR18 secretion from $P_{AOX1}$ | $P_{AOX1}$-$\alpha_{sec}$-SR18-$TT_{AOX1}$ | Zeocin | $P_{AOX1}$ (PmeI) | This study |
| SMH243* | SR18 secretion from decoupled system | $P_{C120}$- $\alpha_{sec}$-SR18-$TT_{AOX1}$ | Zeocin | $P_{AOX1}$ (PmeI) | This study |
| SMH244* | Intracellular Mfp5 from decoupled system | $P_{C120}$-Mfp5-$TT_{AOX1}$ | Zeocin | $P_{AOX1}$ (PmeI) | This study |

* These plasmids also contain $P_{AOX1}$ which is solely used for integration at the $P_{AOX1}$ locus and does not drive expression of any genes. Plasmids highlighted in yellow belong to the toolkit to easily implement light induction of protein production in *K. phaffii*.

Abbreviations: TT – Terminator, $\alpha_{sec}$ – α-factor secretion tag, $\beta_{lacto}$ – β-lactoglobulin

### Supplementary Table 2. *K. phaffii* strains used in this study.

| Strain | Description | Genotype | Plasmid used to construct | Source |
|---|---|---|---|---|
| NRRL Y-11430 | Wild-type parent strain | WT | N/A | ATCC |
| ySMH3-100-3 | Intracellular yEGFP with coupled system at a single copy. | NRRL Y-11430 $P_{AOX1}$::($P_{C120}$-yEGFP-$TT_{AOX1}$,$P_{ADH2}$-EL222-$TT_{ADH2}$, ZeoR) | SMH131 | This study |
| ySMH3-1000-3 | Intracellular yEGFP with coupled system at multiple copies. | NRRL Y-11430 $P_{AOX1}$::($P_{C120}$-yEGFP-$TT_{AOX1}$,$P_{ADH2}$-EL222-$TT_{ADH2}$, ZeoR) | SMH131 | This study |
| ySMH5-100-5 | Intracellular yEGFP from $P_{AOX1}$ at a single copy. | NRRL Y-11430 $P_{AOX1}$::($P_{AOX1}$-yEGFP-$TT_{AOX1}$, ZeoR) | SMH5 | This study |
| ySMH5-1000-3 | Intracellular yEGFP from $P_{AOX1}$ at multiple copies. | NRRL Y-11430 $P_{AOX1}$::($P_{AOX1}$-yEGFP-$TT_{AOX1}$, ZeoR) | SMH5 | This study |
| ySMH29 | *his4* knockout strain | NRRL Y-11430 *his4* ($P_{HTX1}$-Cas9-$TT_{DAS1}$, $P_{HTX1}$-gRNA$_{HIS4}$-$TT_{AOX1}$) | SMH60 | This study |
| ySMH51-D2 | β-lactoglobulin secretion by $P_{AOX1}$ (top producer from screen) | NRRL Y-11430 $P_{AOX1}$::($P_{AOX1}$-$\alpha_{sec}$-$\beta_{lacto}$-$TT_{AOX1}$, ZeoR) | SMH71 | This study |
| ySMH115-14 | $P_{C120}$-yEGFP to characterize decoupled system | NRRL Y-11430 $P_{AOX1}$::($P_{C120}$-yEGFP-$TT_{AOX1}$, ZeoR) | SMH6 | This study |
| ySMH154-1 | yEGFP secretion from $P_{AOX1}$ (intermediate strain for verifying phototoxicity in secretion strains) | NRRL Y-11430 $P_{AOX1}$::($P_{AOX1}$-$\alpha_{sec}$-yEGFP-$TT_{AOX1}$, ZeoR) | SMH158 | This study |
| ySMH154-C5 | yEGFP secretion from $P_{AOX1}$ (top producer from screen) | NRRL Y-11430 $P_{AOX1}$::($P_{AOX1}$-$\alpha_{sec}$-yEGFP-$TT_{AOX1}$, ZeoR) | SMH158 | This study |
| ySMH185-1 | For verifying phototoxicity in secretion strains | ySMH154-1 $P_{ADH2}$::($P_{ADH2}$-EL222-$TT_{ADH2}$, KanR) | SMH139 | This study |
| ySMH186-9 | Intracellular Mfp5 from $P_{AOX1}$ (top producer from screen) | NRRL Y-11430 $P_{AOX1}$::( $P_{AOX1}$-Mfp5-$TT_{AOX1}$, ZeoR) | SMH207 | This study |
| ySMH193-11 | Intracellular yEGFP with one copy of EL222 (used for decoupled system characterization) | ySMH115-14 $P_{ADH2}$::($P_{ADH2}$-EL222-$TT_{ADH2}$, KanR) | SMH139 | This study |
| ySMH193-6 | Intracellular yEGFP with three copies of EL222 (used for decoupled system characterization) | ySMH115-14 $P_{ADH2}$::($P_{ADH2}$-EL222-$TT_{ADH2}$, KanR) | SMH139 | This study |
| ySMH193-9 | Intracellular yEGFP with eight copies of EL222 (used for decoupled system characterization) | ySMH115-14 $P_{ADH2}$::($P_{ADH2}$-EL222-$TT_{ADH2}$, KanR) | SMH139 | This study |
| ySMH196 | Intermediate strain for ROS scavenger tests (no EL222) | ySMH29 $P_{AOX1}$::($P_{AOX1}$-$\alpha_{sec}$-yEGFP-$TT_{AOX1}$, ZeoR) | SMH158 | This study |
| ySMH197 | Intermediate strain for ROS scavenger tests (has EL222) | ySMH196 $P_{ADH2}$::($P_{ADH2}$-EL222-$TT_{ADH2}$, KanR) | SMH139 | This study |
| ySMH203-16 | Decoupled circuit parent strains (1 copy of EL222) | NRRL Y-11430 $P_{ADH2}$::($P_{ADH2}$-EL222-$TT_{ADH2}$, KanR) | SMH139 | This study |
| ySMH203-8 | Decoupled circuit parent strains (3 copies of EL222) | NRRL Y-11430 $P_{ADH2}$::($P_{ADH2}$-EL222-$TT_{ADH2}$, KanR) | SMH139 | This study |

| | | | | |
|---|---|---|---|---|
| ySMH203-2 | Decoupled circuit parent strains (5 copies of EL222) | NRRL Y-11430 $P_{ADH2}$::($P_{ADH2}$-EL222-$TT_{ADH2}$, KanR) | SMH139 | This study |
| ySMH203-21 | Decoupled circuit parent strains (8 copies of EL222) | NRRL Y-11430 $P_{ADH2}$::($P_{ADH2}$-EL222-$TT_{ADH2}$, KanR) | SMH139 | This study |
| ySMH207 | *CTA1* co-expression strain | ySMH197 *his4*::($P_{GAP}$-*CTA1*- $TT_{AOX1}$, HIS4) | SMH219 | This study |
| ySMH209 | *SOD1* co-expression strain | ySMH197 *his4*::($P_{GAP}$-*SOD1*- $TT_{AOX1}$, HIS4) | SMH229 | This study |
| ySMH210 | *PMP20* co-expression strain | ySMH197 *his4*::($P_{GAP}$-*PMP20*- $TT_{AOX1}$, HIS4) | SMH230 | This study |
| ySMH217-A2 | yEGFP secretion with 1 copy of EL222 (top producer from screen) | ySMH203-16 $P_{AOX1}$::($P_{C120}$- $\alpha_{sec}$-yEGFP-$TT_{AOX1}$, ZeoR) | SMH234 | This study |
| ySMH219-D7 | yEGFP secretion with 5 copies of EL222 (top producer from screen) | ySMH203-2 $P_{AOX1}$::($P_{C120}$- $\alpha_{sec}$-yEGFP-$TT_{AOX1}$, ZeoR) | SMH234 | This study |
| ySMH223-G5 | SR18 secretion from $P_{AOX1}$ (top producer from screen) | NRRL Y-11430 $P_{AOX1}$::($P_{AOX1}$-$\alpha_{sec}$-SR18-$TT_{AOX1}$, ZeoR) | SMH242 | This study |
| ySMH224-A11 | β-lactoglobulin secretion with 1 copy of EL222 (top producer from screen) | ySMH203-16 $P_{AOX1}$::($P_{C120}$- $\alpha_{sec}$-$\beta_{lacto}$-$TT_{AOX1}$, ZeoR) | SMH240 | This study |
| ySMH226-C1 | β -lactoglobulin secretion with 5 copies of EL222 (top producer from screen) | ySMH203-2 $P_{AOX1}$::($P_{C120}$- $\alpha_{sec}$- $\beta_{lacto}$ -$TT_{AOX1}$, ZeoR) | SMH240 | This study |
| ySMH230-A6 | SR18 secretion with 1 copy of EL222 (top producer from screen) | ySMH203-16 $P_{AOX1}$::($P_{C120}$- $\alpha_{sec}$-SR18-$TT_{AOX1}$, ZeoR) | SMH243 | This study |
| ySMH231-G2 | SR18 secretion with 5 copies of EL222 (top producer from screen) | ySMH203-2 $P_{AOX1}$::($P_{C120}$- $\alpha_{sec}$- SR18-$TT_{AOX1}$, ZeoR) | SMH243 | This study |
| ySMH238-9 | Intracellular Mfp5 with 1 copy of EL222 (top producer from screen) | ySMH203-16 $P_{AOX1}$::( $P_{C120}$-Mfp5-$TT_{AOX1}$, ZeoR) | SMH244 | This study |

Abbreviations: ZeoR – Zeocin resistance, AmpR – Ampicillin resistance, KanR – Kanamycin/G418 resistance, $\beta_{lacto}$ – β-lactoglobulin. Strains highlighted in yellow belong to the toolkit to easily implement light induction of protein production in *K. phaffii*.

**Supplementary Table 3. Primers used for qPCR analysis**

| Target | Amplicon length (bp) | Primer | Sequence |
|---|---|---|---|
| *ARG4* | 150 | SMHpP280 | AAGTCCATTCCGTCAACCTATAAC |
| | | SMHpP281 | TAGAGCATTCTTCATTCGTTCG |
| yEGFP | 104 | SMHpP284 | GGTGAAGGTGAAGGTGATGC |
| | | SMHpP285 | CCGAAAGTAGTGACTAAGGTTGG |
| EL222 | 167 | SMHpP420 | TTCGGAGCAGATGATACTAGAG |
| | | SMHpP421 | TCCTCAGAATACCCAGTTAGATC |

**Supplementary Sequence S1. $P_{C120}$ promoter sequence**

TAGGTAGCCTTTAGTCCATGCGTTATAGGTAGCCTTTAGTCCATGCGTTATAGGTAG
CCTTTAGTCCATGCGTTATAGGTAGCCTTTAGTCCATGCGTTATAGGTAGCCTTTAGT
CCATGCTTAAGAGACACTAGAGGGTATATAATGGAAGCTCGACTTCCAG

**Supplementary Sequence S2. EL222 transcription factor sequence.** The full sequence of the EL222 transcription factor with its various regions annotated by color: Kozak – green, SV40 nuclear localization signal – purple, VP16 activation domain – red, EL222 – blue.

TAAAATCATGGCTCCGAAAAAAAAGAGAAAAGTAGCCCCACCAACGGACGTTTCAC
TTGGTGACGAATTACATTTAGACGGTGAGGATGTAGCTATGGCACATGCCGATGCTC
TTGATGATTTTGATCTGGACATGTTAGGTGATGGGGATTCTCCCGGCCCCGGATTTA
CACCACATGATTCTGCTCCGTATGGCGCGTTGGACATGGCAGATTTCGAATTTGAAC
AGATGTTTACTGATGCACTTGGTATCGACGAATATGGAGGTGGAGCAGATGATACTA
GAGTTGAAGTCCAACCTCCAGCCCAATGGGTATTGGATTTGATCGAGGCCTCACCCA
TAGCAAGTGTTGTTAGTGATCCCAGATTGGCTGATAATCCTTTGATTGCTATCAACC
AGGCCTTTACCGATCTAACTGGGTATTCTGAGGAAGAATGTGTTGGTCGTAATTGTA
GATTTTTGGCAGGATCTGGTACTGAACCCTGGCTAACAGATAAGATCCGTCAAGGTG
TTCGTGAGCATAAACCCGTGTTGGTTGAGATTTTGAATTACAAAAAGGATGGTACTC
CTTTTCGTAATGCAGTTTTGGTGGCACCAATCTATGATGACGATGATGAATTACTAT
ACTTCCTTGGTAGCCAAGTGGAAGTGGATGACGATCAACCGAACATGGGCATGGCC
CGTAGAGAACGTGCTGCGGAGATGTTAAAAACTCTTTCACCACGTCAATTAGAAGTC
ACAACTCTGGTTGCATCAGGCCTAAGAAATAAAGAAGTGGCTGCCAGACTGGGTCT
TTCAGAAAAAACTGTCAAAATGCACAGAGGTTTGGTAATGGAAAAATTAAACTTAA
AAACGAGTGCAGATTTAGTTAGAATTGCCGTAGAAGCAGGTATTTAA
